# Structural insight into sodium-dependent bile acid transport by members of the SLC10 family

**DOI:** 10.64898/2026.02.20.706973

**Authors:** Criska Yuanting Li, Aurelien Grob, Leah Repa, Oliver Huxley, Deborah H. Brotherton, Patrick Becker, Ruby Dadzie, Oliver Beckstein, Alexander D. Cameron

**Author notes:** These authors contributed equally (Criska Yuanting Li, Aurelien Grob, Leah Repa).

## Abstract

ASBT and NTCP are sodium-coupled secondary transporters of the SLC10 family that play critical roles in the enterohepatic recycling of bile acids. Secondary transporters generally function through the alternating access mechanism. However, in outward-facing structures of NTCP this mechanism is violated with a pore running through the protein. No such pore is observed in previously reported structures of bacterial ASBT homologues, but these proteins have an additional transmembrane-helix adjacent to the binding site. Here we investigate another homologue of ASBT without this extra helix. We solve the crystal structure of the inward-facing state in multiple conformations and a humanised version in the outward-facing state, with and without bile acid bound. Unlike for NTCP there is no pore through the protein as the exit is blocked by the flexible TM6. Consequently, the presence of the pore is not necessarily common to the family. Molecular dynamics simulations of both bacterial and human proteins highlight that while the bile acid substrate is anchored to residues at the centre of the transporter, lipids from the surrounding membrane interact with the hydrophobic sterol group. Incorporating lipids into the transport mechanism can explain how bile acids modified by large hydrophobic groups can bind and be transported.

## Introduction

Bile acids are necessary for the uptake of lipids and fat-soluble vitamins. They are synthesised in the liver by the breakdown of cholesterol and released to the intestine via the bile duct^1^. Rather than being excreted they are then reabsorbed and transported back to the liver through the portal vein. Critical to this process are several membrane transporters^1,2^ including the Apical Sodium-dependent Bile acid Transporter (ASBT) and the Na^+^-dependent Co-transporting Polypeptide (NTCP) of the SLC10 family. ASBT is located in the ileum^3^. Its critical position has made it of interest to the pharmaceutical industry. Small molecule inhibitors were developed as potential cholesterol lowering drugs and while these showed some efficacy in animal models they have not been pursued^4,5^. Instead, these molecules have been repurposed as drugs to treat bowel and cholestatic liver disorders^6–9^, with elobixibat approved in Japan for chronic constipation and odevixibat and maralixibat approved by the FDA for pruritis^10^. Given that *in-vitro* ASBT has also been shown to transport bile acid-drug conjugates, ASBT has also been suggested in a prodrug strategy^11,12^. NTCP is located on the hepatocyte and is responsible for the uptake of bile acids from the portal vein. While physiologically it is also important in bile acid transport, it is of more interest pharmaceutically as it has been hijacked by the hepatitis B virus as an entry receptor^13,14^.

The first high resolution structural information regarding this family came from crystal structures of two bacterial homologues of the human SLC10 family, from *Neisseria meningitidis* (ASBT_NM_)^15^ and *Yersinia frederiksenii* (ASBT_YF_)^16^. These transporters have 10 transmembrane helices arranged into two domains, referred to as core and panel domains. The core domain is characterised by two discontinuous helices that cross-over to form an X shaped motif. Like human ASBT (hASBT) and human NTCP (hNTCP) these proteins are sodium dependent^15,16^, and in the inward-facing ASBT_NM_ structure sodium ions were observed to bind in the core region near to the cross-over helices, with interactions to residues conserved throughout the sodium-dependent members of the family. Two major conformations of the bacterial proteins were observed, an inward-facing state^15,16^, with a cavity open to the interior of the cell and an outward facing state, with the main difference between them in the relative position of the panel with respect to the core^16^. Although, both transporters were shown to transport bile acids *in vitro*, physiologically they are more likely to transport metabolites^17,18^. More recently structures of mammalian NTCP have been solved by cryo-EM in inward^19,20^ and outward-facing states^19,21–23^. Most members of the human SLC10 family have 9 rather than 10 transmembrane helices due to the absence of the first transmembrane helix of the bacterial transporters. Overall, the mammalian NTCP structures are consistent with the bacterial structures, with slight shifts of the helices of the panel domain to counteract the missing TM1. The big surprise, however, with these structures is that all the outward-facing structures have a pore through the centre of the protein. This violates the alternating access model for transporters, in which the substrate-binding site, should not be accessible from both sides of the membrane simultaneously^24,25^. Density at the interface between the panel and core domains led Liu *et al* to propose a novel mechanism relying on the bile acids themselves sealing the pore to prevent molecules diffusing through^20,22^.

Given that four different groups have published very similar open pore structures, using different antibodies/nanobodies, detergents and nanodiscs, it seems unlikely that the open pore structure is an artefact. It raises the question, though, whether this should be true of hASBT and the other nine transmembrane-helix members of the family. Although hASBT and hNTCP share 37% sequence identity and both transport bile acids they have a different profile of compounds that can be transported, with hNTCP seemingly being more promiscuous^26,27^. The cross-reactivities of the clinically relevant inhibitors of hASBT towards hNTCP have also been recently compared showing that these inhibitors bind hNTCP much less tightly^9^.

To investigate this class of proteins further, we investigated a bacterial homologue of hASBT and hNTCP, which, like the human proteins also only contains 9 transmembrane helices. We solved the crystal structures of the wild type (WT) protein in the inward facing state and of a construct of the protein where ten residues in the substrate cavity had been mutated to be more like hASBT, in the outward-facing state. In all structures the sodium ions were clearly defined in the binding site. We also obtained a structure of the outward-facing state in the presence of a bile acid. Unlike hNTCP the outward-facing state does not contain an open pore, due to the difference in position of TM6. Instead, it is much more similar to AlphaFold2^28^ generated structures of hASBT.

## Results

### Structure of ASBT_LB_ in the inward-facing state

Through GFP screening^29^ of bacterial homologues of hASBT we discovered a protein from *Leptospira biflexa* with 9 predicted TM helices and 32% sequence identity to human ASBT (Supplementary Figure 1) that could be expressed and purified (ASBT_LB_). Given that relative to hASBT the only insertions or deletions in ASBT_LB_ (Supplementary Figure 1) are of single residues in the loop regions between TM4 and TM5 and between TM6 and TM7, this construct was thought to be a good model to investigate for providing further insight into the mechanism of this family. Using stability assays^30,31^, we established that the addition of deoxycholate (DCA) or taurocholate (TCA) stabilised the protein (Supplementary Figure 2), consistent with these bile acids binding to the protein. Crystals were grown under a range of conditions using the lipidic cubic phase (LCP) method^32^. The best diffracting crystals resulted in a structure with two molecules in the asymmetric unit (denoted as ASBT_LBWT-Xtal1A_ and ASBT_LBWT-Xtal1B_ respectively) that was refined at 2.2 Å (Supplementary Table 1). The nine transmembrane helices are arranged with TMs 1, 5 and 6 forming the panel domain and TMs 2,3,4,7,8,9 the core domain (Figure1A) similar to ASBT_NM_ (Figure 2a) and hNTCP (Figure 3a). There is a clear cavity extending into the protein from the inward-facing side (Figure 1B).

**Figure 1:**
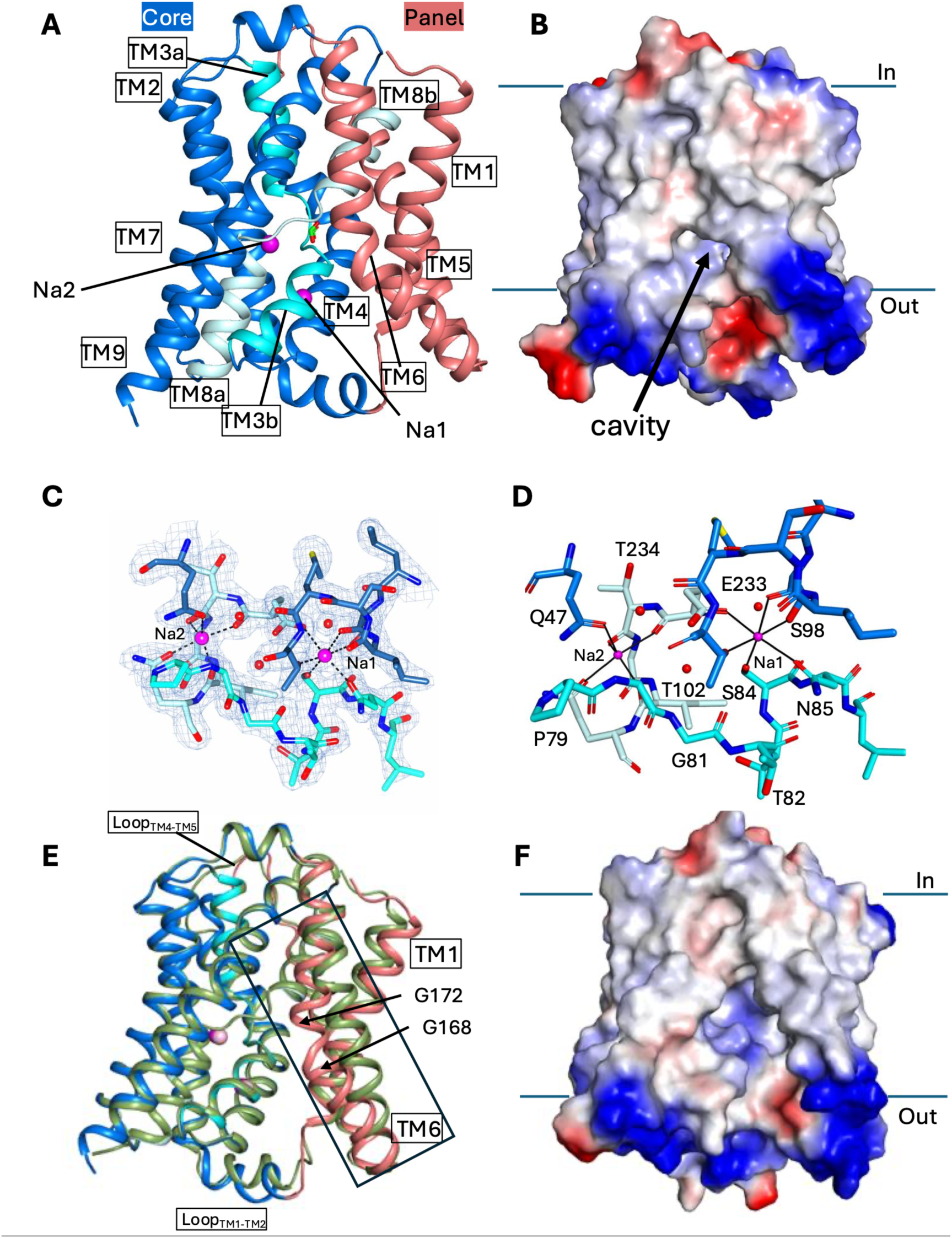
Structure of wild-type ASBT_LB_ in the inward-facing form. **A)** Ribbon representation of ASBT _LBWT-Xtal1A_. The core domain is shown in blue, with the cross-over helices, TM3 and TM8 in light blue and cyan respectively. The sodium ions are shown as magenta spheres and the formate in a stick representation. **B)** Electrostatic representation showing the cavity accessible from the inward-facing side of the protein. **C)** Density associated with the sodium binding site. The stick representation is coloured as in A. The 2mFo-DFc is shown at 1.5σ. **D)** The sodium binding site, coloured as in A. The coordination of the sodium ions is shown by black lines. Water molecules are represented as red spheres. **E)** Superposition of ASBT_LBWT-Xtal2_ (green) on molecule B from ASBT_LBWT-Xtal1A_ (coloured as in A) showing the conformational change to TM6 (boxed). **F)** Electrostatic surface of the ASBT_LBWT-Xtal2_ highlighting the more open cavity.

**Figure 2:**
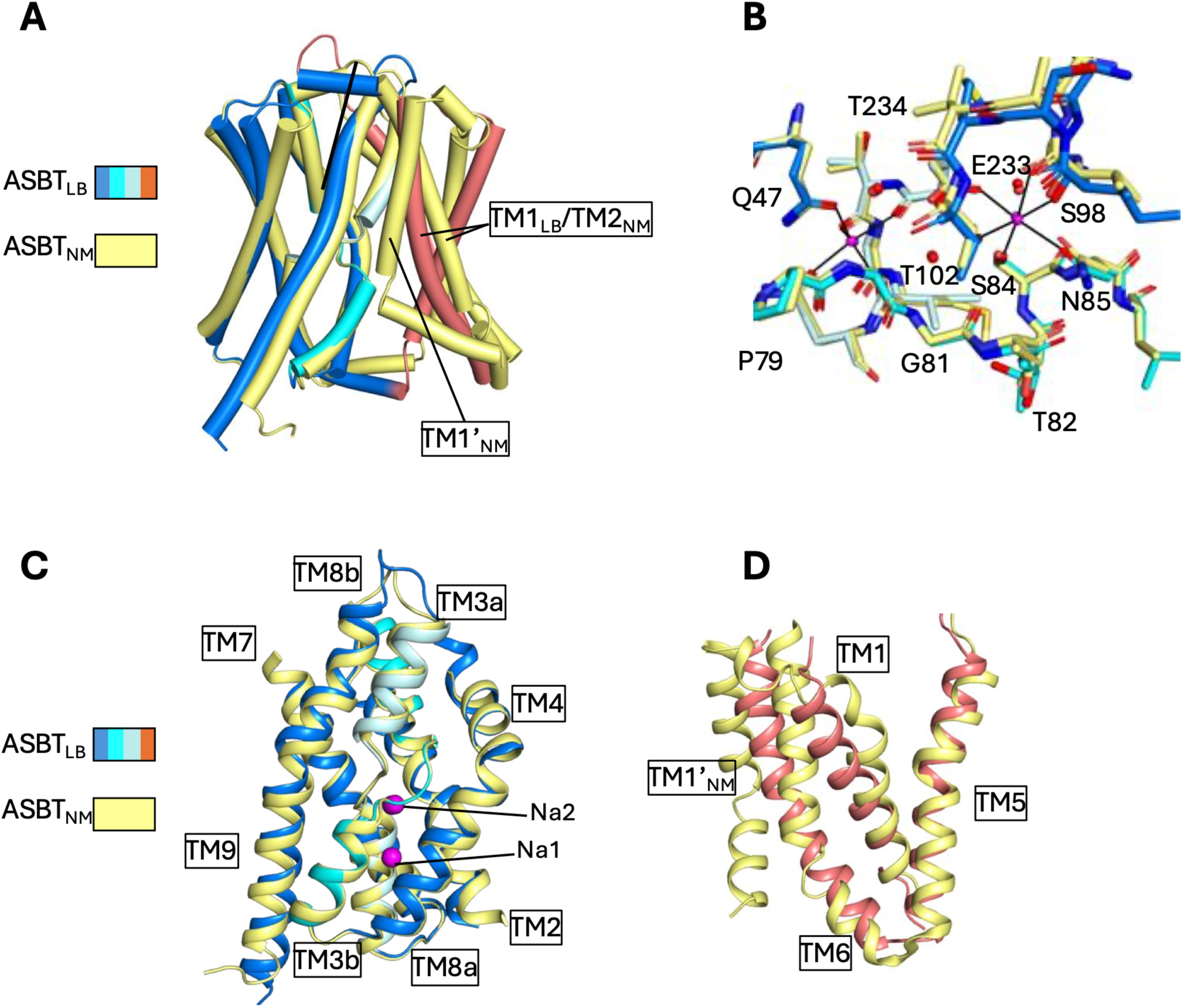
Comparison of ASBT_LB_ to ASBT_NM_. **A)** Overall superposition. ASBT_LB_ is coloured as in Figure 1. ASBT_NM_ is depicted with yellow carbon atoms. **B)** Sodium binding site. **C)** Superposition of core domains. **D)** Overlay of panel domains, following superposition on the core domains of the respective proteins.

**Figure 3:**
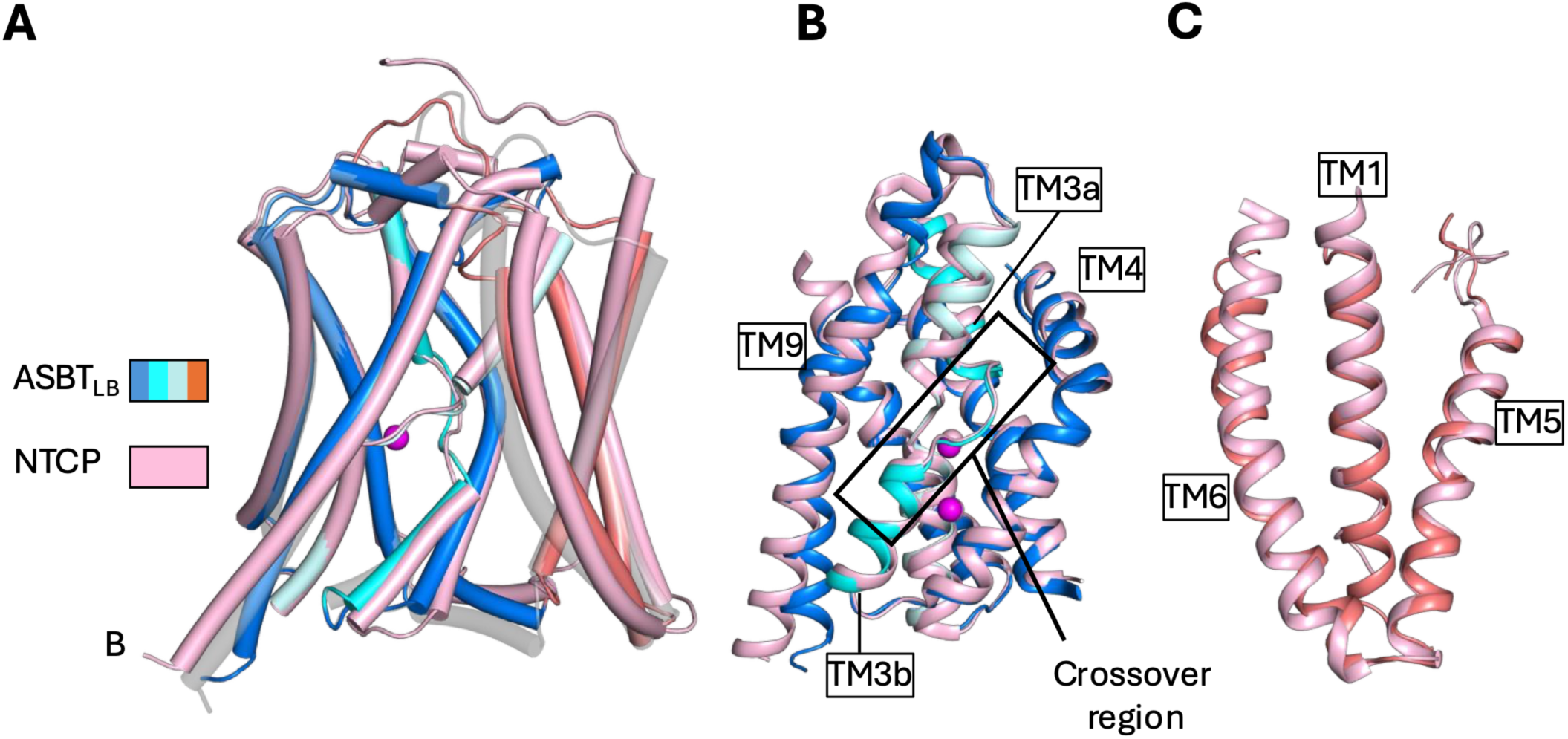
Comparison of ASBT_LB_ to inward-facing hNTCP. **A)** Overall superposition. ASBT_LBWT-Xtal2_ is coloured as in Figure 1a. hNTCP (7PQG) is coloured in pink. A molecule from the ASBT_LBWT-Xtal1_ is shown with partial transparency to illustrate the difference in the position of TM6. **B)** Superposition of the respective core domains with the sodium-bound inward-facing structure (9QZQ) coloured pink including the Na ions. The crossover region is very similar in this structure whereas it differs slightly in the lower resolution hNTCP structure without Na bound (A). (**C)** Superposition of the respective panel domains.

In the core domain there is very clear density present in the expected sodium binding sites (Figure 1C). Na1 is coordinated in an octahedral arrangement by the side chains of S84, N85, S98, T102 and E233 and the main chain carbonyl oxygen of S98 (Figure 1D). Na2 is located at the C-terminus of TM8a (Figure 1A) and is coordinated in a square pyramidal arrangement by the side chains of Q47 and Q237 and the main chain carbonyl oxygens of E233, T234, I236 (Figure 1D). The position of the 6^th^ ligand that would complete the octagonal sphere is in proximity to Gly80 of the linker between TM3a and b (Figure 1D). These residues are conserved amongst the SLC10 family members (Supplementary Figure 3). and the coordination is very similar to that seen in ASBT_NM_ (Figure 2A), which has 26% sequence identity with ASBT_LB_ (Supplementary Figure 3). The water molecules between the two sodium ions seen in ASBT_NM_^15,18^ are also observed in ASBT_LB_ (Fig 1C, Fig 2B). Indeed, the overall conformation of the core domain of ASBT_NM_ is very similar to that of ASBT_LBWT_ with an rmsd of 1.1 Å over 177 atoms within 3.8 Å (Figure 2C). Although DCA was present in the crystallisation mixture, there is no evidence of this bile acid in the density. Instead, in the cross-over region there is density that has been assigned to a formate present in the crystallisation mixture (Supplementary Figure 4A). The overall conformation of the core domain is also similar to that of hNTCP (Figure 3B) with the sodium ions positioned in a similar location to those observed in a recently published inward-facing structure^20^.

In the panel domain more variation is seen amongst the structures. Between the two molecules of the asymmetric unit of ASBT_LB_, there is a slight difference in TM6, that appears to be caused by the presence of a lipid-like molecule in one molecule (Supplementary Figure 4B), which is not present in the other (Supplementary Figure 4A). A second data set from another crystal form was also collected (ASBT_LBWT-Xtal2_). In this structure, which was refined at 2.9 Å and is also in an inward-facing conformation, there is a more pronounced shift in TM6 with the helix flexing around Gly168 and Gly172 to increase the lateral entrance to the cavity (Figure 1E, Supplementary Movie 1). Density consistent with a lipid tail is again present in the cleft between the panel and core domains (Supplementary figure 4C). The shift in TM6 is accompanied by a change in the neighbouring TM1. The only other significant differences between the structures are in the loops connecting TMs 1 and 2 and 4 and 5 between the panel and core domains (Supplementary Movie 1). Overall, the panel domain is more similar to hNTCP than it is to ASBT_NM_, as might be expected given the additional helix in ASBT_NM_, even though the relative position of the panel to the core domain is very similar to all the reported inward-facing structures (Figure 2A and 3A).

### Structure of ASBT_LB_ with selected humanising mutations in the outward-facing state

In order to increase the sequence identity between ASBT_LB_ and hASBT we introduced 10 mutations (ASBT_LB10mut_). The mutations were selected on a combination of sequence conservation, previously reported mutational analysis^33–39^ and the position within the expected binding site (V14L; T83A, T206P, I236M, G239T, P240Q, I243S, A266S, V269Q, P270L, Supplementary Figures 3 and 5). As for the wild-type protein, bile acids were observed to stabilise the ASBT_LB10mut_ protein (Supplementary Figure 2). The protein was crystallised using the LCP method as before. The structure was solved and refined at a resolution of 2.6 Å (Supplementary Table 1). Unlike the structures of the wild-type protein, the protein was clearly in an outward-facing state with the exit to the inward-facing side blocked and a crevice present on the outward-facing side (Figure 4A). Again, although the protein was crystallised with DCA, there was no evidence of this molecule in the binding site. With respect to the wild type structures the panel domain is rotated approximately 22° with respect to the core domain around an axis approximately parallel to TM7, combined with a 5 Å translation along this axis (Figure 4B, Supplementary Movie 1). Overall, the core domains of the inward and outward-facing states are very similar (Figures 4C). As observed for the inward-facing state, sodium ions are clearly present in the binding sites, with a very similar coordination geometry (Figure 4D, Supplementary Figure 5).

**Figure 4:**
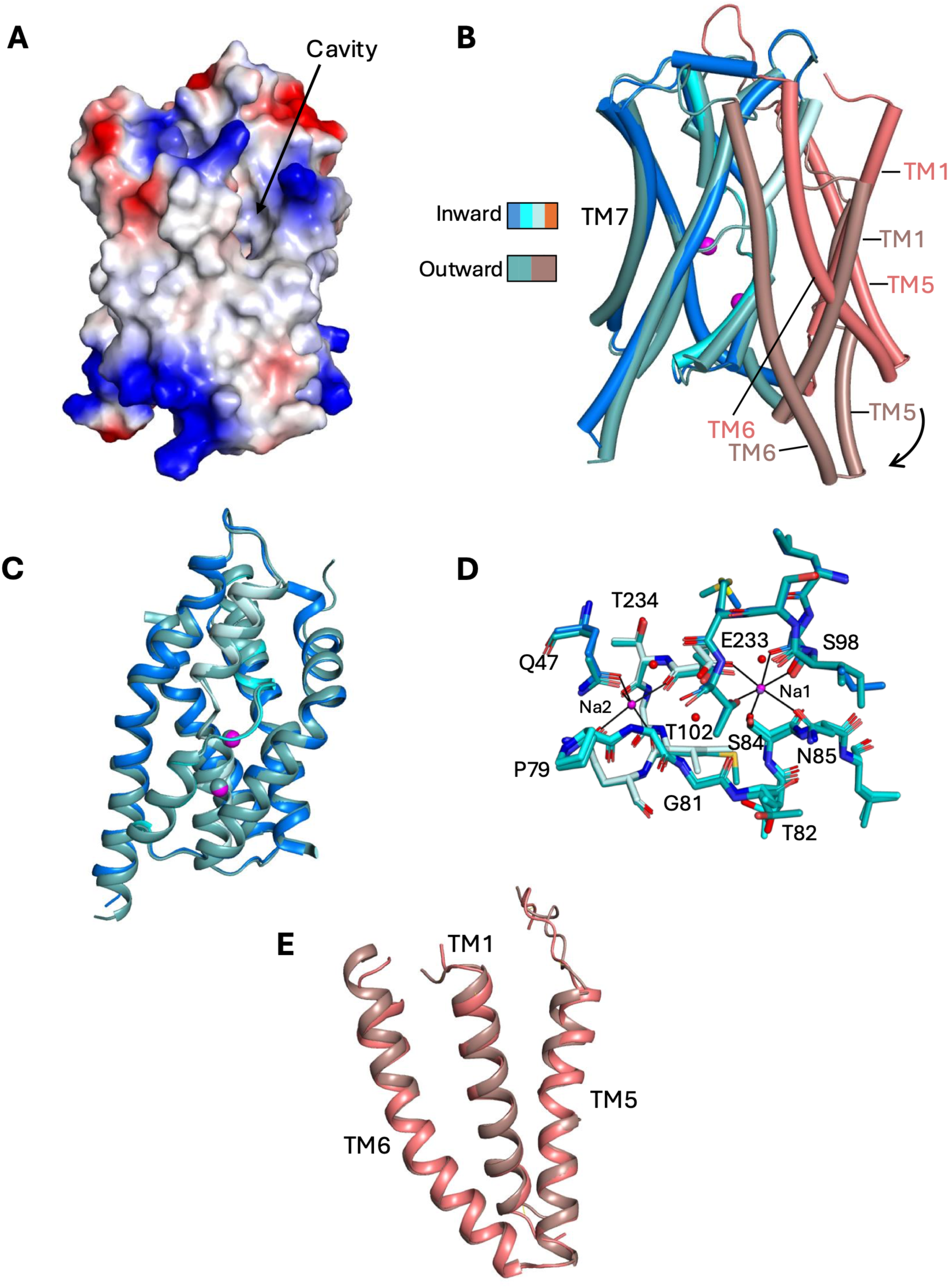
Outward-facing form of ASBT_LB_. **A)** Electrostatic surface illustrating the cavity on the outward-facing side. **B)** Superposition of the outward-facing structure on ASBT_LBxtal2_. The outward-facing structure has been coloured with the core domain in sea green and the panel domain in brown. The two structures have been superposed based on their respective core domains and the arrow indicates the rotation of the panel domain against the core. **C)** Superposition of the core domains. **D)** Superposition of the sodium ion binding site. **E)** Superposition of the panel domains.

The conformation of the panel domain is similar to that of the ASBT_LBWT-Xtal2_, in which TM6 is straighter (Figure 4E). The orientation of the panel with respect to the core domains is very similar to both outward-facing structures of hNTCP (Figure 5A) and ASBT_YF_ (Supplementary Figure 6). With respect to the outward-facing structures of hNTCP, TM6 is much nearer the core domain and contributes to sealing the inward-facing side of the protein (Figure 5A). Consequently, the pore that is observed in hNTCP is not present in ASBT_LB_ (Supplementary Figure 6).

**Figure 5:**
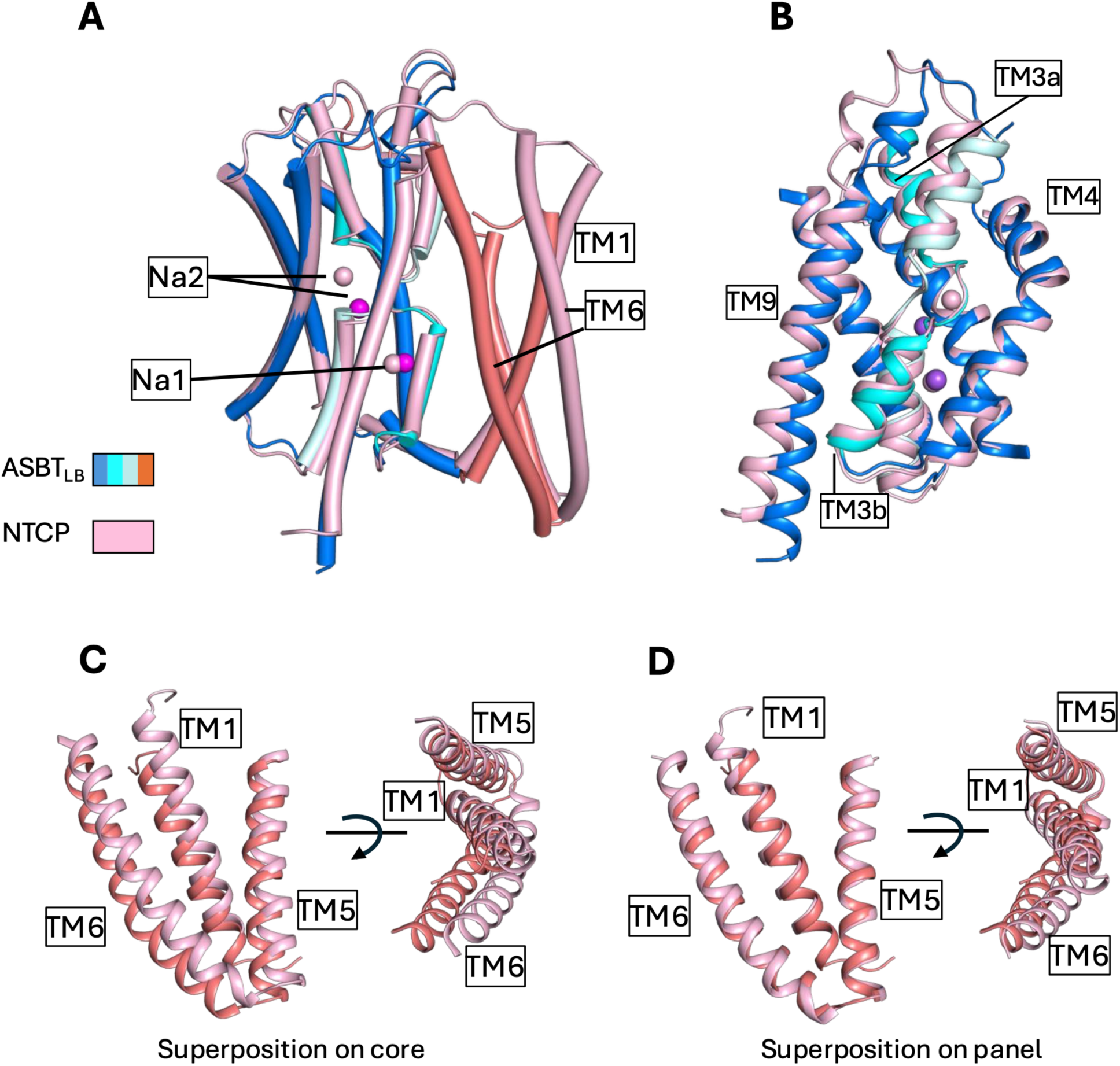
Comparison of outward-facing form of ASBT_LB_ to that of hNTCP. **A)** Overall superposition. ASBT_LB10mut_ is coloured as in Figure 1. hNTCP (7ZYI) is coloured light pink. The Na ions are represented by spheres. **B)** Superposition of the core domains. **C)** Overlay of the panel domains after superposition on the core. **D)** Overlay of the panel domains after superposition on the panel.

### Structure of ASBT_LB10mut_ with deoxycholic acid

Given that no DCA was observed in the binding site with the above data set we then increased the concentration of DCA in the crystallisation solution by adding the DCA to the well solution in addition to the protein. Crystallisation of ASBT_LB10mut_ resulted in 5 data sets from different conditions, all with the same crystal form with the protein in the outward-facing conformation as above. The highest resolution data set was refined at a resolution of 2.5 Å (ASBT_LB10mut-DCA_ Supplementary Table 1). Overall, the conformation of the protein is similar to the previous structure of the same mutant, with the protein in the outward-facing state and bound to sodium ions (Figure 6A). However, very clear density was evident for the deoxycholic acid inserted in the outward-facing side between TM1 and TM5 with the acidic group in proximity to the cross-over region of the core region of the protein (Figure 6B). There are hydrogen bonds to the main chain nitrogen and hydroxyl oxygen of Thr82 (Figure 6C). The hydroxyl oxygen at position 12 of the deoxycholate also interacts with a water molecule, which is within hydrogen bonding distance of Ser124 and Asn238. Overall, there is very little conformational change between DCA bound and free protein structures except a slight twisting of the N-terminus of TM1 and a shift of the linker regions between the 2 domains following TM6 (Figure 6A, Supplementary Movie 1). In proximity to the carboxylic acid group we see density that we have not been able to assign unambiguously. In the final structure water molecules have been placed in this density. The position of the DCA is different from the positions of the glycochenodeoxycholic acid or taurocholic acid that have been modelled for hNTCP^22,40^ (Figure 6E). Structures of hNTCP have also been solved as a complex with bulevirtide, a peptide designed as an HBV entry inhibitor^41,42^. The position of DCA, overlaps with that of bulevirtide in hNTCP (Figure 6F).

**Figure 6:**
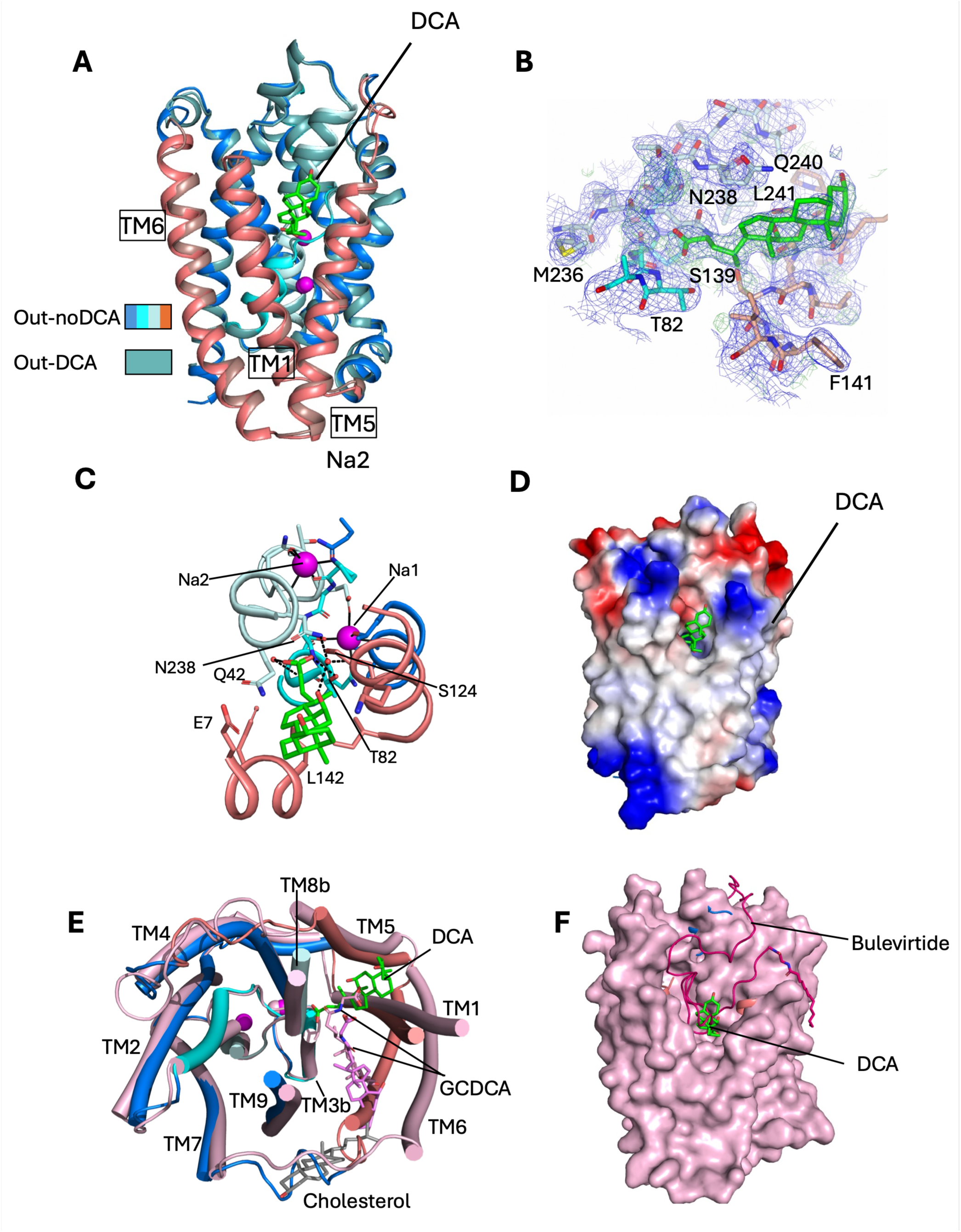
Binding of deoxycholate to the outward-facing ASBT_LB_. **A)** Superposition of the DCA-bound structure on the non-bound structure. The non-bound structure is depicted as in Figure 5. The DCA-bound structure is coloured with the core domain in teal and the panel domain in brown. The DCA is depicted in sticks with green carbon atoms. **B)** Electron density associated with the DCA. The 2mFo-DFc (blue) and mFo-DFc (green) have been calculated based on the structure before the DCA was added and the final refined structure has been superposed. **C)** The binding site of the DCA and the cross-over region of the protein. Potential hydrogen bonding interactions are shown with black dashed lines, whereas the sodium coordination is depicted by solid black lines. **D)** Electrostatic surface representation. **E)** Superposition on the outward-facing hNTCP structure with glyochenodeoxycholate (GCDCA) modelled in the structure (7ZYI). The GCDCA is depicted with pink sticks. A cholesterol from the same structure is shown with grey carbon atoms. **F)** The structure of ASBT_LB10mut_ complexed with DCA superimposed on the structure of hNTCP complexed with the peptide bulevirtide (8rqf). hNTCP is shown as a surface representation in pink with the bulevirtide as a cartoon in dark red. ASBT_LB10mut_ is shown as a cartoon with the DCA in a stick representation with green carbon atoms. Given the similarity between the hNTCP and the ASBT_LBmut10_ the latter is mostly hidden.

### Comparison with AlphaFold models of hASBT

As there was no experimental structure of hASBT we used AlphaFold2 to predict inward and outward-facing hASBT structures. The highest rank inward-facing structure that we calculated had an rmsd of 1 Å for 281 out of 288 pruned atom pairs, relative to the wild-type ASBT_LBXtal1A_ structure with alpha helices well-matched (Supplementary Figure 6B). TM6 in the predicted outward-facing structures of hASBT was more similar to the position of TM6 in ASBT_LB10mut_ as opposed to hNTCP. Comparatively, AlphaFold2 predictions for hNTCP revealed TM6 to be more open than observed in experimental cryo-EM structures (Supplementary Figure 7E). Consequently, unlike hNTCP, the predicted outward-facing structure of hASBT is closed at the inward-facing side as was seen for ASBT_LB10mut_ and does not show the same pore seen in hNTCP.

### Molecular dynamics

To investigate substrate binding further, we performed multiple all-atom MD simulations of ASBT_LB_ (WT) and ASBT_LB10mut_ in outward facing (OF) and inward facing conformations (IF). To generate an IF model of ASBT_LB10mut_ we applied all mutations to the IF WT structure; to create an OF model for WT we reversed all 10 mutations in the OF ASBT_LB10mut_ structure. As a baseline, we ran one simulation of the WT apo protein in OF and IF and these remained stable with no notable conformational changes, reflected in a backbone RMSD below 2 Å over 1 µs (Supplementary Figure 7A). The fully loaded transporter contains two Na^+^ ions and one bound DCA. For the OF conformation, we used the DCA-bound OF crystal structure as the starting conformation. For the IF structure, we used the apo WT IF crystal structure as a starting point, and, in the absence of an experimental IF structure with DCA bound, we employed flexible docking with AutodockVina^43^ to generate a binding pose for WT and then transferred the binding pose to ASBT_LB10mut_ as described in Methods. WT OF with DCA bound was generated by reversing the 10 mutations in the ASBT_LB10mut_ DCA-bound structure. We validated our docking protocol with the OF structure where we recovered the experimental binding pose to within 1.5 Å substrate RMSD (data not shown). In triplicate repeat simulations (duration 1 µs each) the protein was stable with backbone RMSDs below 2.5 Å (Supplementary Figure 7B for ASBT_LB10mut_ and Supplementary Figure 7D for WT), indicative of simulations of high-quality structures in stable conformational states. In the following we focus on the ASBT_LB10mut_ simulations. In the OF conformation, DCA maintains the binding pose observed in the crystal structure with the carboxylate group near Thr82 and Ala83 and the sterol moiety pointing towards the extracellular side (Figure 7A). During the simulations, the DCA binding site and the occupied sodium binding sites Na2 and Na1 are only accessible from the outside as indicated by the water density (Supplementary Figure 8A). In the inward facing conformation, the sterol moiety of DCA switched its orientation to point towards the intracellular side while the carboxylate remains anchored to Thr82 and Ala83 (Figure 7B). Access to the binding site switched to the intracellular compartment, as indicated by the water density that reaches all the way towards Na1 to Na2 and Thr82 and Ala83 (Supplementary Figure 8B). A detailed analysis of the interactions of DCA with the protein showed clearly that in both OF and IF conformation, the carboxylate group forms a hydrogen bond to Thr82 and the backbone carbonyl of Ala83 (Figure 7C). Hydrophobic interactions differ almost completely between OF (eight unique interactions) and IF (nine unique interactions), with only T82 and Q240 appearing for both OF and IF (Figure 7D). For the ASBT_LB10mut_ simulations the binding interactions are much better defined for OF as seen from the RMSD of DCA after superposition on the protein (Supplementary Figure 7C) where RMSDs for OF remain under 4 Å while IF RMSDs rise up to 10 Å. Nevertheless, DCA remains in the binding site and in one simulation, recovers the initial interactions. The WT simulations with DCA bound show even more clearly that Thr82 (and Thr83, corresponding to Ala83 in the mutant) hold the carboxylate of DCA in place in both OF and IF conformation (Supplementary Figure 9A) while the hydrophobic interactions differ (Supplementary Figure 9B). In the WT IF simulations, DCA generally remains closer to its initial binding pose (which was obtained from docking) whereas the OF simulations show larger rearrangements (Supplementary Figure 7E). The comparatively larger deviations from the initial DCA binding pose occurred with introduction of mutations to either create the WT OF DCA-bound structure or the ASBT_LB10mut_ IF DCA-bound structure.

**Figure 7:**
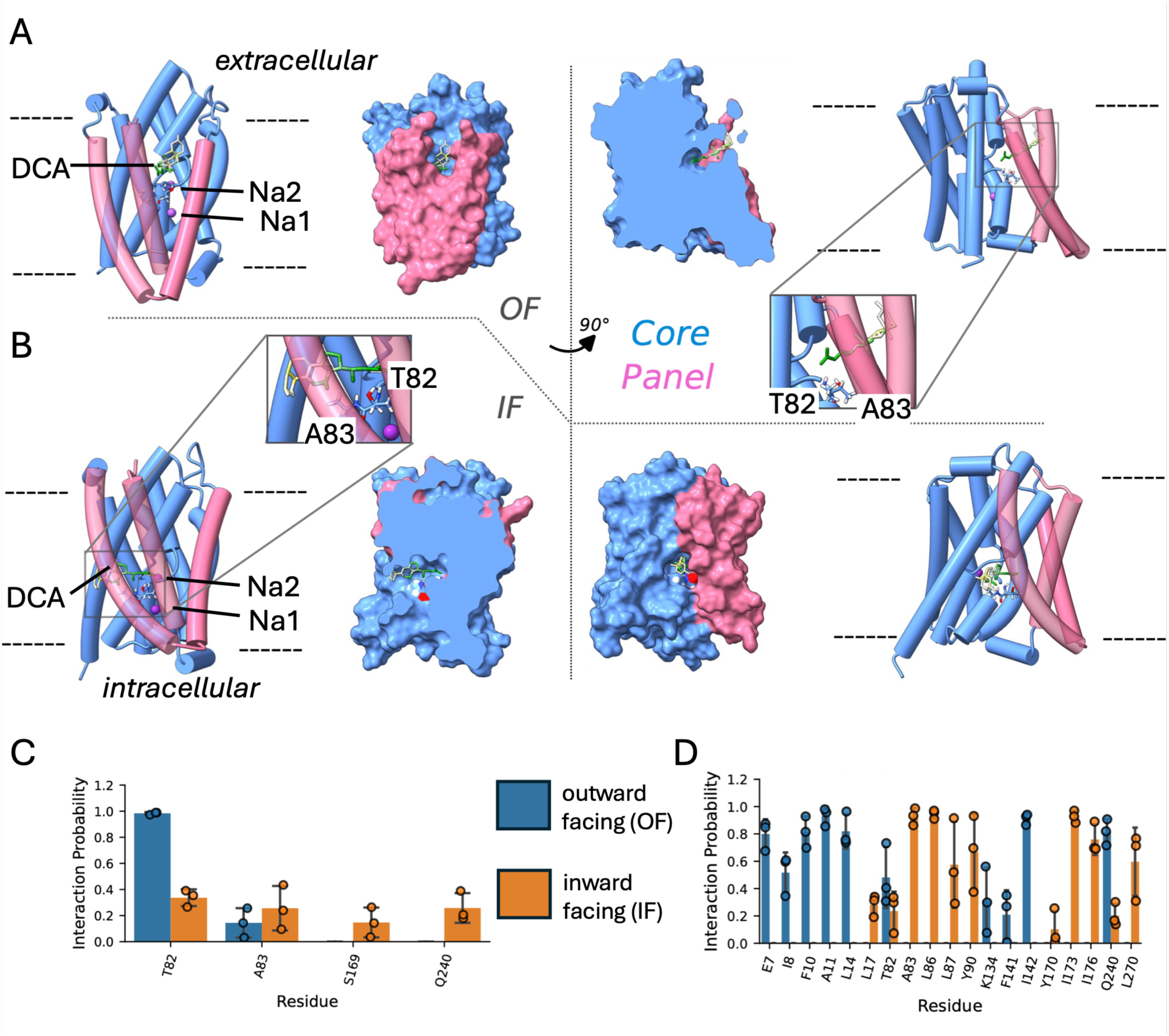
MD simulations of the 2 Na^+^/1 DCA loaded transporter in OF and IF conformation. **A)** Structural diagram of ASBT_LB10mut_ and DCA in OF conformation in both cartoon and VDW density representations, with core and panel domains (blue and pink), DCA (white, yellow and green atomic representations), residues T82 and A83 (labelled in inset), and Na^+^ ions (purple spheres) shown. **B)** IF conformation. **C)** Probability of hydrogen-bond acceptor interaction between DCA and ASBT_LB10mut_ in OF (blue) and IF (orange) conformation. **D**) Hydrophobic contacts interaction probability. Results from three independent MD simulations in either OF or IF conformation (points) were averaged (bars). Error bars indicate the standard deviation over the three replicates.

**Figure 8:**
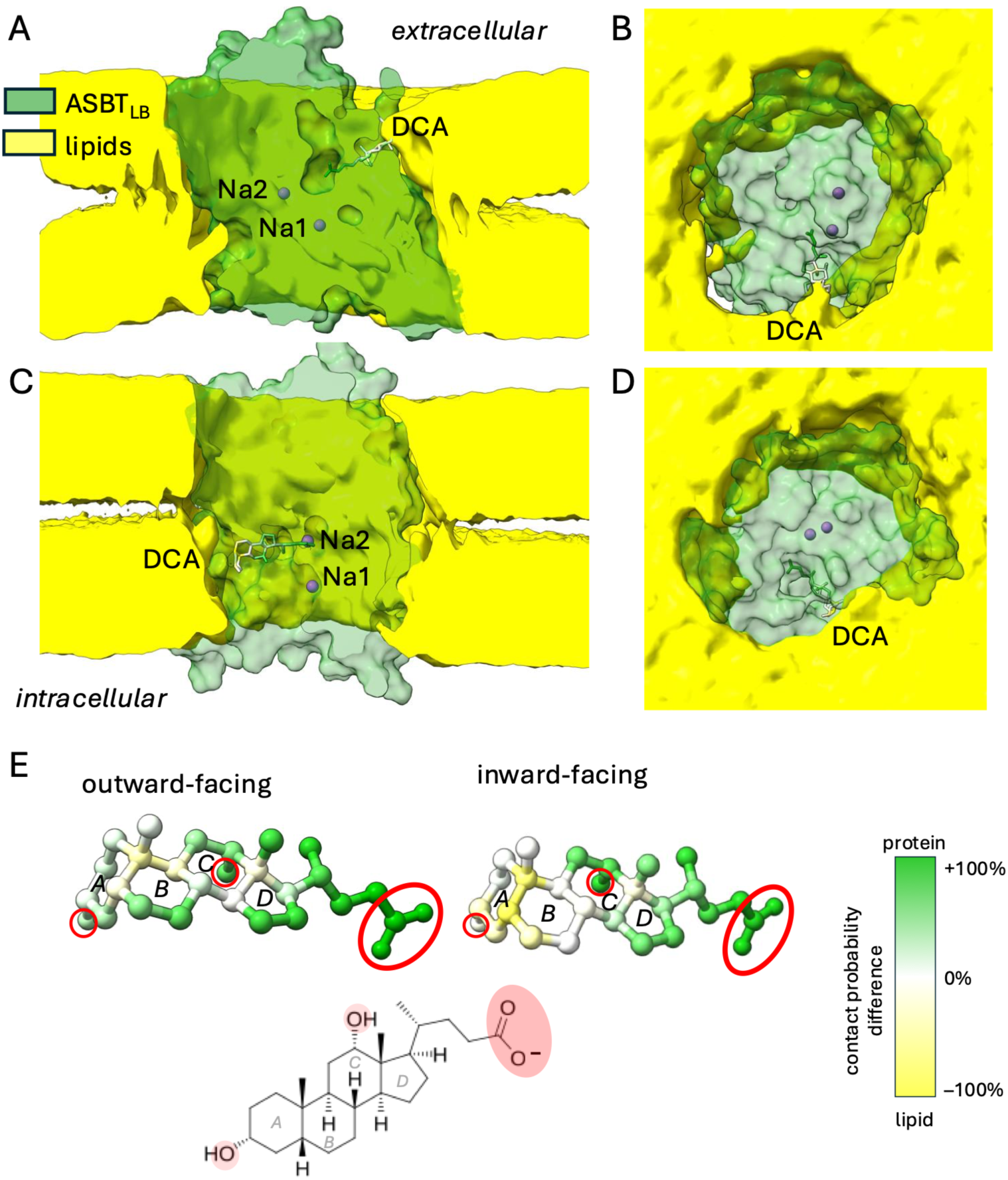
Proteo-lipidic substrate binding from MD simulations. Density representations showing ASBT_LB10mut_ (green) embedded in lipid membrane (yellow), along with DCA (white, green and yellow atomic representation) and Na^+^ ions (purple spheres). Note that ASBT_LB10mut_ is slightly transparent to show the location of Na^+^ ions within. DCA is coloured according to calculated protein/lipid interactions, with colour corresponding to interaction partner (green for protein, yellow for lipids). **A)** Sideview of the OF conformation. **B**) View on the extracellular side of the transporter in the OF conformation, showing the accessibility of the DCA binding site. **C)** Side view of the IF conformation. **D)** View on the intracellular side of the transporter in the IF conformation with a visible pathway towards the DCA binding site. **E)** Interaction of DCA from outward-facing (left) and inward-facing (right) MD simulations with lipids and proteins together with chemical structure (bottom) with polar groups highlighted in red and rings labelled *A*–*D*. Hydrogen atoms are omitted for clarity from the molecular images. Each atom in the DCA conformer is coloured by the difference in its interaction probability with any protein or any lipid atom. Green (+100%) indicates prevalent interaction with the protein while yellow (–100%) indicates exclusive interaction with lipids.

DCA also interacts with the lipids of the membrane during the simulations. The binding site is not fully contained inside the protein so that the lipids of the membrane contribute to binding in both OF and IF conformations (Figures 9A-D). During the simulations the hydrophilic carboxylate group remained tethered to Thr83 while the hydrophobic sterol moiety (rings A-D in Figure 9E) formed contacts more frequently with lipids than with the protein. Especially when bound to the OF conformation of ASBT_LB_, the first three steroid rings (A, B, and C) are primarily interacting with membrane lipids as the proteinaceous binding site opens up sideways to the membrane. In the IF conformation, our predicted binding pose is situated more deeply in the protein itself although the water-accessible pathway is partially confined by the membrane lipids, thus forming a proteo-lipidic pore (Figure 9C,D and Supplementary Figure 8B). The simulations with WT confirm these results, as shown in Supplementary Figure 9C and 8C,D.

We also investigated hASBT using MD. The AlphaFold2 predicted structure was unstable in simulations and could not be used (data not shown). However, given that, with the exception of the N and C termini, there were only two single residue insertions or deletions between hASBT and ASBT_LB_ we modelled hASBT into the electron density of the inward-facing ASBT_LB_ and the outward-facing ASBT_LB10mut_. This resulted in very few clashes and MD simulations of both the outward and inward-facing states were stable with backbone RMSDs <2.5 Å (Supplementary Figure 7F); these models performed remarkably well and would have been difficult to distinguish from actual crystallographic models based on the simulations, unlike the AlphaFold2 model, which was unusable. The OF model represents an outward-facing structure as indicated by the water accessibility of the DCA and the occupied Na1/Na2 binding sites (Supplementary Figure 9E); the IF model appears to be partially occluded (Supplementary Figure 8F). When DCA was modelled into the OF hASBT structure based on the ASBT_LBmut10-DCA_ crystal structure in the outward-facing state, DCA maintained its initial binding pose for ∼300ns before adopting a slightly different conformation (Supplementary Figure 7G). Binding to the IF conformation was similar to the ASBT_LBmut10_ simulations. The hASBT simulations reproduce the major findings from the simulations of the bacterial structures: Residue Thr110 (corresponding to Thr82) binds DCA via its carboxylate in both IF and OF conformation (Supplementary Figure 9D) while the sterol moiety encounters completely different protein environments (Supplementary Figure 9E) and partially interacts directly with the lipids of the membrane (Supplementary Figure 9F).

## Discussion

The canonical alternating access mechanism of membrane transport relies on the substrate binding site of a transporter being accessible in turn from either one side of the membrane or the other without ever opening a solvent-accessible channel through the protein^24,25,44^. There are three main mechanisms that have been proposed: rocking bundle, rocker switch and elevator mechanism^45^. The first two involve the protein rotating around an axis in the centre of the protein to alternately switch the protein from one conformation to the other. With the elevator mechanism, on the other hand, one domain appears to lift with respect to the other^45^. Following the structure elucidation of ASBT_YF_ in outward and inward-facing conformations, the SLC10 proteins were proposed to follow the elevator mechanism^16^, similar to the structure of Na^+^/H^+^ antiporters, which have a similar fold^46^. With the publication of structures of hNTCP by various groups an apparent variation of the mechanism was observed: although two conformations of the protein were observed in line with the bacterial proteins^19^, the outward-facing structures have an open pore with access to the binding site from either side of the membrane^19,21–23^. No other experimental structure has been reported of a mammalian SLC10 family member. Here we solve the structure of a bacterial homologue of the SLC10 proteins with nine transmembrane helices, mimicking hASBT and hNTCP. To make it more hASBT-like we also introduced 10 mutations in areas where substrate is likely to bind, based on the extensive studies of hASBT^33–39^. Interestingly, while the wild-type protein crystallised exclusively in the inward-facing state, all the structures we obtained of the mutant are outward-facing. This suggests that the introduced mutations push the equilibrium between the inward and outward-facing states towards the latter. Overall, the conformations of ASBT_LB_ in the inward and outward-facing states are very similar to those of the corresponding structures of hNTCP^19^. Unlike hNTCP, however, due to the difference in position of TM6, the open pore that is observed in hNTCP is sealed in ASBT_LB_.

ASBT is sodium-coupled^47,48^. According to the conventional alternating access model, the protein can switch conformations either when both substrate and sodium ions are present or when neither are there in order to avoid dissipating the sodium gradient^44^. The positions of the sodium ions were clearly defined in the ASBT_NM_ inward-facing structure and have been suggested to rigidify the structure of the cross-over region of the protein, which is likely to be important in substrate binding^15,18^. For ASBT_LB_ we observe the coordination of the sodium ions to be very similar to ASBT_NM_ in both outward and inward-facing states, and in MD simulations, two Na^+^ ions modelled into the Na1 and Na2 binding sites remain stably bound over 1 µs. In hASBT the equivalent mutation of any of these residues causes complete loss of transport activity^34,37,38,49^, with similar results in hNTCP^50^. While in hNTCP, the coordination of the sodium ions is similar in the inward-facing form^20^, in the only structure of outward hNTCP with sodium ions modelled^22^, the position of Na1 mimics ASBT_LB_, even though Na2 is displaced by 5.5 Å and is not as well coordinated by the protein^22^.

The density that we observe in the binding site is unambiguously DCA, having the characteristic shape of the molecule with an approximately 90° angle between the first and second rings of the cholate. This arrangement places the acidic group near to the cross-over region of the protein as would be expected of a substrate that would be transported in a sodium-dependent manner^18^. However, as located in our structure, steric clashes would presumably prevent transport with the classical elevator mechanism and raises the question how DCA will move during transport. While one mechanism proposed for NTCP involves transition of the bile acid though a pore, for both ASBT_YF_ and NTCP a more conventional mechanism has been suggested necessitating some movement of the bile acid^16,51^. This involves flipping of the substrate as observed for the omega-3 fatty acid transporter from the MSF family^52^ and would be consistent with our structures. ASBT transports conjugated forms of bile acids, such as taurocholate and glycocholate. Furthermore, *in vitro* drug conjugates of bile acids are able to be transported, suggesting that even bigger molecules can bind^53^. The additional density that we observe in the binding site that we have not been able to assign could easily be replaced by these larger moieties. Given that sodium binding is equivalent in both the outward and the inward-facing states it seems that the sodium ion sets up the substrate binding site. For ASBT_NM_ it has been suggested that binding causes a change to Thr112 in the cross-over region^18^. Here we see Thr82, the equivalent residue, interacting with the DCA, but don’t see a significant change in the residues of the cross-over region. The MD simulations of the outward facing and inward facing conformation suggest that Thr82 and the neighbouring Ala83 act as anchors for the carboxylate of the deoxycholate substrate while the hydrophobic sterol moiety is partially enclosed by the lipid membrane. There is emerging evidence of the membrane lipids being involved in transport pathways for membrane proteins. For ion channels that enable the movement of lipids across the membrane, such as the TMEM16 family of lipid scramblases^54–56^, the polar headgroup of the lipids moves through the protein and the acyl chains extend into the surrounding membrane. Similarly, for the OSCA/TMEM63 mechanosensitive channels^57^, a proteo-lipidic pore was described as the ion pathway. Our MD simulations indicate that a hydrophobic part of DCA interacts with lipids and the lipid membrane and is apparently integral for accommodating and thus binding the substrate. Although the architecture of ASBT_LB_ protein would not be compatible with opening a channel as seen in the TMEM16 and 63 ion channels, it is tempting to speculate that the membrane environment provides unspecific binding for bulky hydrophobic moieties while the protein specifically requires a long carboxylate tail to anchor the polar end. In the absence of experimental structures of the intermediate steps of transport or MD simulations of the full substrate translocation process we cannot say whether the lipid is involved in transport as well as binding.

The question must be asked how relevant the structures we present here are to the human bile acid transporters. The conformations of the inward and outward-facing states of hNTCP are very similar to ASBT_LB_, except for TM6. However, this helix shows the most flexibility in both ASBT_LB_ and hNTCP^19,51^. In ASBT_LB_, the helix flexes around two glycine residues (Figure 1E and Supplementary Movie 1). While only one of these glycine residues is conserved in hNTCP, both are conserved in hASBT (Supplementary Figure 3). In the inward-facing form of ASBT_LB_, the conformation of TM6 appears to be correlated to the presence of lipid like molecules between panel and core domains. It is in this region that bile acids have been modelled in the outward-facing open pore conformation of hNTCP. However, an analysis of the cryo-EM maps from the various hNTCP structures, shows similar density in this region, independently of whether or not the bile acids were reported to be added during purification and vitrification (see eg EMD-31837) and the density is not sufficiently clear to unambiguously assign it to being bound bile acids. Rather than the open pore structure that is observed in outwards-facing structures of hNTCP, in our structures, TM6 packs onto the core domain to form a much more closed state. It is possible that a similar conformation could exists in hASBT, as using AlphaFold2^28,58^ to predict the structure we do not observe any structure with an open pore and MD simulations don’t show any large changes. This is not the same in AlphaFold2 predicted structures of hNTCP where there is a large space between the two; notably, molecular dynamics simulations provide evidence that hNTCP can adopt an outward-facing structure with a closed pore^51^.

### Overall transport mechanism

Taken together our data support an alternating access mechanism of transport with the energetic barrier for transitions between IF and OF state being small. The mechanism doesn’t necessarily involve opening a pore through the protein as seen for hNTCP, though the flexibility of TM6 would mean that a pore could be opened transiently. The sodium ions will shape the binding site by stabilising the conformation of the cross-over region of the protein. The substrate can then interact with the residues of the cross-over region, with the sterol moiety of the substrate interacting with the surrounding lipids. Such a proteo-lipidic interaction may explain how substrates with bulky hydrophobic groups can bind and consequently be transported.

## Materials and Methods

### Expression and Purification

A Blast search was carried out for bacterial homologues of hASBT predicted to contain 9 transmembrane helices. 6 sequences were selected (Uniprot entries: B0SJT7, N1W7E1, A0A1Y5H5S3, A0A2A6RHM8, A0A3E0WHV7, A0A1T5HQR4). The genes encoding the sequences were codon optimised for expression in *E. coli* and purchased as gBlocks (Integrated DNA Technologies). These were inserted into a modified version of the expression vector, pWaldo GFPd^29^ in which the TEV protease site had been altered to a site for recognition by 3C protease^59^. GFP fusions were expressed in *E. coli* Lemo21 (DE3) cells following the MemStar^60^ procedure and screened for expression and suitability for expression using Fluorescence Size Exclusion Chromatography (FSEC) ^61^. A protein from *Leptospira Biflexa* (Uniprot B0SJT7) was selected for further study. Cultures were grown at 37 °C, 200 rpm, in PASM-5052 media supplemented with 0.25 mM rhamnose. When cultures reached an OD_600_ of 0.5, the temperature was dropped to 25 °C and 0.4 mM IPTG was added for protein induction overnight.

Cell pellets were harvested by centrifugation and resuspended in PBS (137 mM NaCl, 2.7 mM KCl, 10 mM Na_2_HPO_4_, 1.8 mM KH_2_PO_4_) with 1 mM MgCl_2_, DNaseI, and 0.5 mM 4-benzenesulfonyl fluoride hydrochloride (AEBSF) and disrupted by passing three times through a cell disruptor at 25 kPsi. Cell lysate was centrifuged at 24,000 g at 4 °C for 12 minutes to remove insoluble cell debris, and the supernatant was subjected to ultracentrifugation at 200,000 g, 4 °C for 45 minutes. Membrane pellets were resuspended in PBS, 15 mL per 1 l of culture, snap frozen in liquid nitrogen, and then stored at -80 °C.

For crystallisation membranes from 3 L of culture were solubilised in 1x PBS pH 7.4, 150 mM NaCl, 1% DDM for 2h at 4°C and ultracentrifuged for 45 mins, 4 °C, 200,000g to remove insoluble material. Imidazole was added to 10 mM, and the membrane suspension was mixed with 1 ml of Ni-NTA Superflow resin (Qiagen) per 1mg of GFP–His8 and incubated for 3 hours or overnight at 4 °C. Slurry was decanted into a glass Econo-Column (Bio-Rad) and washed with typically, 10 Column Volumes (CV) of 1x PBS (PH7.4), 150 mM NaCl, 10 mM imidazole, 0.03% DDM, 20 CV of the same buffer but with 20mM imidazole, 15 CV of 20mM Tris pH 7.5, 150mM NaCl, 30mM Imidazole pH 7.5, 0.03% DDM. The 3C-GFP-His8 tag was then cleaved overnight by addition of equimolar amounts of 3C protease directly to the protein-bound resin. The following day, the cleaved protein FT was collected and chased through with 5CV of the final wash buffer. The eluted protein was pooled and passed over a 5ml Ni-NTA His-Trap column (GE Healthcare). Protein was concentrated to 8-10mg/ml in a concentrator (Sartorius) with at 100K cut off and loaded onto a Superdex 200 Increase 10/300 GL gel filtration column (Cytiva) equilibrated in 20mM Tris pH 7.5, 150mM NaCl, 0.03% DDM. Samples containing ASBT_LB_ were collected and concentrated to 25-30mg/ml.

To create ASBT_LBmut10_ the gene encoding the protein was purchased as a codon optimised gBlock (Integrated DNA Technologies) and was expressed and purified as above, except that the concentration of rhamnose was reduced to 0.1mM rhamnose during expression.

### Stability Assay

To test how bile acids affected the stability of the protein an assay based on the binding of 7-Diethylamino-3-(4’-Maleimidylphenyl)-4-Methylcoumarin (CPM) to the protein^30,31^ was used. CPM (ThermoFisher) was dissolved in 100% DMSO to a final concentration of 4 mg/ml. The assay was performed in 0.2 ml non-skirted low profile 96-well PCR plates (ThermoFisher). 2.5 μg of protein in 20 mM Tris-HCl (pH 7.5), 150 mM NaCl, 0.03% DDM was added to each well supplemented with the concentration of each compound of interest varying from 10μM to 1.5mM. The 4mg/ml CPM dye was diluted 1:540 in 20 mM Tris-HCl (pH 7.5), 150 mM NaCl, 0.03% DDM and 17 μl of the diluted dye was added to each well. The assay was performed using a Stratagene Mx3005P Real-Time PCR machine (Strategene) with an emission wavelength of 463 nm and excitation wavelength of 387 nm, and samples were heated from 25°C to 95°C in 1°C/min steps. Data were analyzed using GraphPad Prism.

### Crystallisation and Structure solution

#### ASBT_LB-xtal1_

Protein at a concentration of 25-30mg/ml was subjected to crystallisation using the lipidic cubic phase method of crystallisation^32^. The protein with 1mM DCA was mixed with monoolein at a 60:40 (w/w) ratio using a coupled syringe device (Molecular Dimensions Ltd UK) and crystallisation trials were set up at 20°C using glass sandwich plates with a Mosquito Robot. An initial X-ray diffraction data set extending to 3.5Å was obtained from crystals grown in 0.2M Lithium chloride, 0.1M Sodium acetate pH 4.5, 42% PEG 400. The crystals were cryo-cooled in liquid nitrogen and data were collected at beamline I24 at Diamond Light Source, UK. Data were processed in DIALS^62^. The structure was solved with Phaser^63^ using the structure of ASBT_NM_ (3ZUY) with its first helix omitted as the search model, before preliminary refinement was carried out in Phenix Liebschner, 2019 #2480} and Refmac^64^ interspersed with manual rebuilding with Coot^65^. A higher resolution data set (ASBT_LB-xtal1_) was subsequently collected from crystals grown in 0.12M Magnesium formate dihydrate, 0.1M Sodium chloride, 0.1M Tris pH 8.5, 33% PEG 600. Data were collected at beamline I24 at Diamond Light Source, UK. Despite the diffraction pattern showing multiple crystals the data were successfully processed using the automated processing pipelines at Diamond Light Source. Essentially the data were processed in DIALS^62^ through the Xia2 pipeline^66^. The structure of the protein was solved using the DIMPLE^67^ pipeline at Diamond with the preliminary structure from the lower resolution data set. Due to the overlapping diffraction patterns of multiple crystals in the loop, the merging statistics were poorer than would be expected, however, the electron density was consistent with a 2.2Å data set as given by the automated processing. Further refinement was carried out in Phenix^68^ interspersed with manual rebuilding using Coot^65^.

#### ASBT_LB-xtal2_

Protein was crystallised as above but without the addition of DCA. Data were collected from crystals grown in 0.32M Lithium chloride 0.1M Sodium citrate pH 5.5, 14% w/vPEG 4000 from condition F4 of Memgold2 (Molecular Dimensions). Prior to freezing the crystals were soaked for a short time in a solution containing 1mM TCA. The data were processed with DIALS^62^ through the Xia2 pipeline and scaled in Aimless^69^ in the CCP4 suite^70^. Molecular replacement was carried out in Phaser^63^ using ASBT_LB-xtal1a_ as the search model^62^ and refinement carried out as above.

#### ASBT_LBmut10_

Protein was crystallised as above with the addition of 1mM DCA. Data were collected from a crystal grown in sodium chloride 0.1M Tris 8.5 24% v/v PEG 350 MME from Memgoldmeso C7 (Molecular Dimensions). The data were processed with DIALS^62^ through the Xia2 pipeline and scaled in Aimless^69^ in the CCP4 suite^70^. Molecular replacement was carried out in Phaser using ASBT_LB-xtal1a_ as the search model^63^ with the core and panel domains treated as separate search models. The model and maps were improved using the Buccaneer software^71^ before being refined as above.

#### ASBT_LBmut10-DCA_

Protein was crystallised above with the addition of DCA also being added to the well solutions to a final concentration of 1mM. Data were collected from a crystal grown in 0.12M Magnesium formate dihydrate 0.1M Sodium chloride, 0.1M Tris, pH 8.5, 33% v/v PEG 600, supplemented with 1 mM DCA. The data were processed with DIALS Waterman, 2016 #2476} through the Xia2 pipeline and then scaled in Aimless^69^ in the CCP4 suite^70^. Molecular replacement was carried out in Phaser^63^ using ASBT_LB10mut_ as the search model with refinement carried out as above.

### Modelling human ASBT and NTCP

#### AlphaFold

AlphaFold models for outward and inward-facing models were made in AlphaFold2^28^ through ColabFold^58^ setting the max_msa to 16:32 and the number of seeds to 8. Of the 40 models produced the highest ranking outward and inward-facing structures were chosen.

#### Using crystallographic models as templates

A model of inward-facing hASBT was created in Coot ^65^ by mutating individual residues of the inward-facing ASBT_LBWT-Xtal1B_, choosing the best rotamer to avoid any clashes and using the real-space refine option in Coot to refine against the associated map. The model of the outward-facing hASBT was made in the same way using the structure of ASBT_LB10mut_ as the reference model and map.

Superpositions were performed in ChimeraX^72^

### Docking with Autodock Vina

Autodock Vina^43^ was used to generate candidate poses for DCA binding to inward facing ASBT. Flexible protein-ligand docking was performed on IF ASBT_LB_ using the DCA structure captured in OF ASBT_LB10mut_, with a search exhaustiveness of 2048 within a cell of 2nm x 2nm x 5nm centred on the geometric centre of the protein. The most favourable binding position identified received a Vina score of -8.76 and was used as the starting binding pose for DCA in all IF ASBT_LB_ simulations. This binding pose was also used in all IF ASBT_LB10mut_ and hASBT simulations. The DCA binding poses for these modified protein structures were obtained by aligning these models with the WT IF ASBT_LB_ structure upon their respective protein backbones and using the resulting DCA binding pose relative to the modified structure. The second and third most favourable poses generated by Autodock Vina, with scores of -7.63 and -7.12 respectively, were not used but are shown in comparison to the best DCA pose in Supplementary Figure 11.

### System setup with CHARMM-GUI

All systems were prepared for simulation as a protein-membrane system using CHARMM-GUI’s Membrane Builder tool^73–75^. The protein structures were embedded in a 4:1 palmitoyloleoylphosphatidylethanolamine:palmitoyloleoylphosphatidylglycerol (POPE:POPG) lipid bilayer, with positioning within the membrane solved via PPM 2.0^76^. The protein-membrane system was then surrounded by a water box at pH 7 in the +Z and -Z directions (above and below the membrane), simulating the extra- and intracellular solutions respectively. NaCl was added at a concentration of 150 mM to the solutions on either side of the membrane, with additional Na^+^ or Cl^-^ ions added as needed to neutralize the total charge of the system. The initial box size and total number of atoms generated for each simulation are listed in Supplementary Table 2.

CHARMM-GUI’s GROMACS input generator^77^ was used to generate simulation and forcefield files. All simulations were prepared as an *NPT* system at a temperature of 300 K and pressure of 1 atm, parameterized with the CHARMM36 all-atom forcefield^78^, with the TIP3P water model and CHARMM36 parameters for lipids^79^. DCA was parametrized with the CHARMM General Force Field^80^ (CGenFF) but required manual fixing for incorrect stereochemistry.

### MD simulations with GROMACS

All MD simulations were performed as all-atom equilibrium NPT simulations using GROMACS 2023.3^81^. Simulations were prepared for production runs following the standard CHARMM-GUI protocol generated for *NPT* GROMACS simulations, with an initial energy minimization run followed by multiple equilibration runs with a total pre-production simulation time of ∼1.1 ns. Production runs of apo ASBT_LB_ in both conformations were performed once, while all other production simulations (those that included a bound DCA and two sodium ions in the Na1 and Na2 sites) were performed in three repeats, each prepared from the same set of pre-production simulations but generated uniquely. All production runs were carried out with a 2 fs timestep for a total simulation time of 1 μs each.

These simulations used the v-rescale^82^ thermostat and the, c-rescale^83^ barostat. Long range electrostatics were calculated with the smooth particle-mesh Ewald (SPME) method^84^ while van der Waals forces were switched smoothly to zero between 1.0 nm and 1.2 nm as implemented in GROMACS. The Verlet neighbour list was updated dynamically by GROMACS. The P-LINCS algorithm^85^ was used to constrain all bonds involving hydrogen atoms, except water molecules for which the analytical SETTLE algorithm^86^ was used.

### Data Analysis

MDAnalysis^87^ was used for post-processing of MD trajectories and the calculation of RMSD values as well as protein-DCA and lipid-DCA contact data. The proLIF library^88^ was used to generate protein-DCA interaction fingerprints, following the default protein-ligand procedure. These interaction fingerprints provide a binary metric on whether a particular interaction is occurring between DCA and a given protein residue on each simulation frame. This data was then used to calculate interaction probabilities by dividing the number of simulation frames with detected interactions by the total number of frames recorded.

### Visualisation

Structural images were prepared in PyMol^89^, ChimeraX^72^ and VMD^90^. Images involving electron density were prepared in CCP4mg ^91^. The movie was prepared in ChimeraX.

## Acknowledgements

We thank the staff at beamline I24 at DLS and the technical support in the School of Life Sciences, University of Warwick. The project was supported by a BBSRC MIBTP studentship to AG (BB/M01116X/1). PB was supported by National Institutes of Health (Award No R01GM118772) to OB. DHB was supported by MRC (MR/P010393/1) and Leverhulme Trust (RPG-2022-323).

## Author contributions

The project was initiated by AC and supervised by AC and OB. Cloning, expression, purification, crystallisation and assays were carried out by AG, CL, OH, RD, PB, with supervision by DB. Data collection, processing and structural analysis were carried out by AG, CL and AC. Simulations were carried out by LR and analyzed by LR and OB. AC wrote the manuscript with contributions from all authors.

## Data and materials availability

Data and coordinates have been deposited in the RCSB Protein Data Bank under accession numbers 28SV (ASBT_LB-xtal1_), 28SW ASBT_LB-xtal2_), 28SX (ASBT_LB10mut_) and 28SZ (ASBT_LB10mutdca_). MD trajectories in the GROMACS XTC format were deposited in the OSF.io repository under DOI 10.17605/OSF.IO/8RXZC under the open CC-BY Attribution 4.0 International license.

