## Supplementary Tables and Figures for "Structural insight into sodium-dependent bile acid transport by members of the SLC10 family"

### Supplementary Information

**Supplementary Table 1. Data collection and refinement statistics**

| | $ASBT_{LB\text{-}Xtal1}$ | $ASBT_{LB\text{-}Xtal2}$ | $ASBT_{LB10mut}$ | $ASBT_{LB10mut\text{-}DCA}$ |
|---|---|---|---|---|
| Wavelength | 0.999 | 0.619 | 0.619 | 0.619 |
| Resolution range | 56 - 2.2 (2.28 - 2.21) | 47 - 2.9 (3.68 - 2.92) | 32 - 2.6 (2.85 - 2.59) | 59 - 2.52 (2.72 - 2.52) |
| Space group | P $2_1$ $2_1$ $2_1$ | I 1 2 1 | C 2 2 $2_1$ | P $2_1$ $2_1$ $2_1$ |
| Unit cell | 65.6 94.9 107.6 90 90 90 | 62.5 67.1 75.2 90 91.04 90 | 81.8 110.1 74.4 90 90 90 | 55.2 75.1 95.0 90 90 90 |
| Total reflections | 213123 (14547) | 41189 (20669) | 86775 (20997) | 138627 (24933) |
| Unique reflections | 34241 (2770) | 6831 (3389) | 10742 (2620) | 13781 (2641) |
| Multiplicity | 6.2 (5.3) | 6.0 (6.1) | 8.1 (8.0) | 10.1 (9.4) |
| Completeness (%) | 99.8 (98.8) | 99.71 (99.68) | 99.20 (98.26) | 99.23 (98.32) |
| Mean I/sigma(I) | 6.13 (0.94) | 3.99 (2.14) | 4.91 (1.31) | 4.14 (0.64) |
| Wilson B-factor | 33.00 | 45.39 | 36.75 | 50.23 |
| R-merge | 0.2404 (1.621) | 0.3478 (0.788) | 0.2812 (1.203) | 0.3455 (2.539) |
| R-meas | 0.264 (1.804) | 0.3812 (0.8624) | 0.2999 (1.286) | 0.3632 (2.681) |
| R-pim | 0.1069 (0.7837) | 0.1537 (0.3457) | 0.1031 (0.4486) | 0.1097 (0.8434) |
| CC1/2 | 0.466 (0.403) | 0.972 (0.805) | 0.99 (0.837) | 0.995 (0.606) |
| CC* | 0.797 (0.758) | 0.993 (0.944) | 0.998 (0.955) | 0.999 (0.869) |
| Reflections used in refinement | 34186 (2751) | 6819 (3385) | 10677 (2598) | 13695 (2634) |
| Reflections used for R-free | 1658 (119) | 329 (162) | 539 (134) | 668 (128) |
| R-work | 0.1775 (0.2374) | 0.2400 (0.2632) | 0.2555 (0.3114) | 0.2809 (0.3809) |
| R-free | 0.2256 (0.2882) | 0.2952 (0.3313) | 0.2861 (0.3414) | 0.2927 (0.4044) |
| Number of non-hydrogen atoms | 4956 | 2229 | 2290 | 2271 |
| Macromolecules | 4484 | 2227 | 2257 | 2234 |
| ligands | 379 | 2 | 30 | 36 |
| solvent | 93 | 0 | 3 | 1 |
| Protein residues | 576 | 286 | 289 | 286 |
| RMS(bonds) | 0.009 | 0.009 | 0.002 | 0.002 |

| | | | | |
|---|---|---|---|---|
| RMS(angles) | 1.05 | 1.19 | 0.56 | 0.53 |
| Ramachandran favored (%) | 97.38 | 96.13 | 98.26 | 93.97 |
| Ramachandran allowed (%) | 2.62 | 2.82 | 1.74 | 5.32 |
| Ramachandran outliers (%) | 0.00 | 1.06 | 0.00 | 0.71 |
| Rotamer outliers (%) | 1.59 | 0.00 | 1.18 | 1.99 |
| Clashscore | 3.81 | 7.62 | 2.33 | 5.63 |
| Average B-factor | 39.61 | 43.24 | 55.73 | 59.51 |
| macromolecules | 38.12 | 43.26 | 55.65 | 59.59 |
| ligands | 57.24 | 30.43 | 63.50 | 55.17 |
| solvent | 39.36 | | 36.78 | 42.29 |

Statistics for the highest-resolution shell are shown in parentheses.

**Supplementary Table 2: MD simulations.**

| Run ID | Conformation / Model | Ligands | Initial system size (nm), *X* x *Y* x *Z* | Num. atoms | Repeats x sim. time |
|---|---|---|---|---|---|
| LB-OF-APO | OF / LB WT | None | 9.5 x 9.5 x 11.0 | 95592 | 1 x 1μs |
| LB-IF-APO | IF / LB WT | None | 10.1 x 10.1 x 10.9 | 106544 | 1 x 1μs |
| LB-OF-DCA | OF / LB WT | 2 $Na^+$, 1 DCA | 10.1 x 10.1 x 10.4 | 101532 | 3 x 1μs |
| LB-IF-DCA | IF / LB WT | 2 $Na^+$, 1 DCA | 10.1 x 10.1 x 10.8 | 105812 | 3 x 1μs |
| 10MUT-OF-DCA | OF / $LB_{10mut}$ | 2 $Na^+$, 1 DCA | 10.1 x 10.1 x 10.4 | 102095 | 3 x 1μs |
| 10MUT-IF-DCA | IF / $LB_{10mut}$ | 2 $Na^+$, 1 DCA | 10.1 x 10.1 x 10.7 | 104455 | 3 x 1μs |
| hASBT-OF-DCA | OF / hASBT | 2 $Na^+$, 1 DCA | 10.1 x 10.1 x 10.8 | 105209 | 3 x 1μs |
| hASBT-IF-DCA | IF / hASBT | 2 $Na^+$, 1 DCA | 10.1 x 10.1 x 10.4 | 100944 | 3 x 1μs |

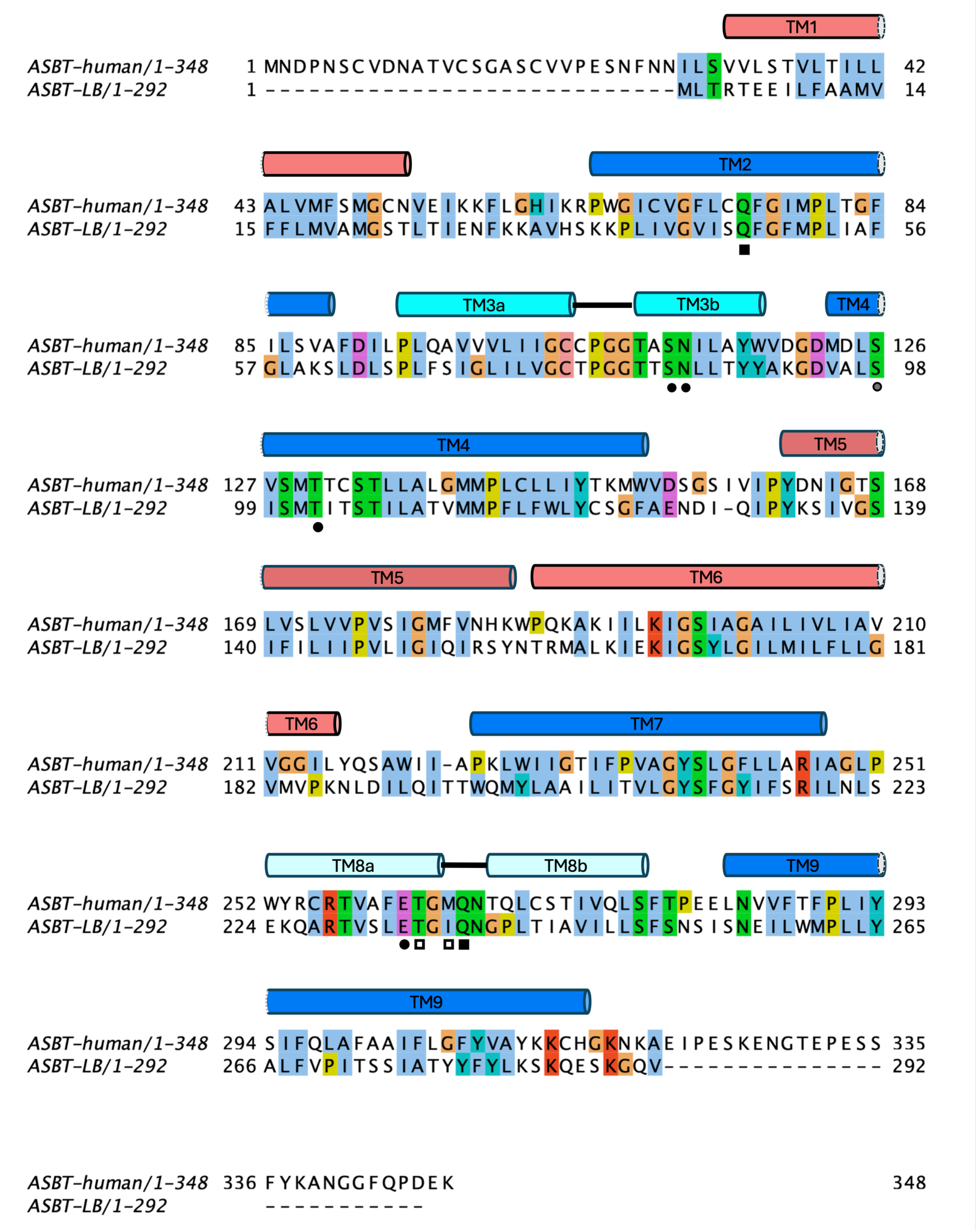


**Supplementary Figure 1: Sequence alignment of ASBT$_{LB}$ and hASBT**. The secondary structure of ASBT$_{LB}$ is shown coloured as in Figure 1. Residues coordinating Na1 by side chains are shown as filled circles and through the main chain as open circles with grey depicting both. Residues coordinating to Na2 are similarly depicted by squares. With respect to hASBT (Uniprot Q12908) ASBT$_{LB}$ has a single residue insertion in the loop between TM4 and TM5 and a single residue deletion in the loop between TM6 and TM7.

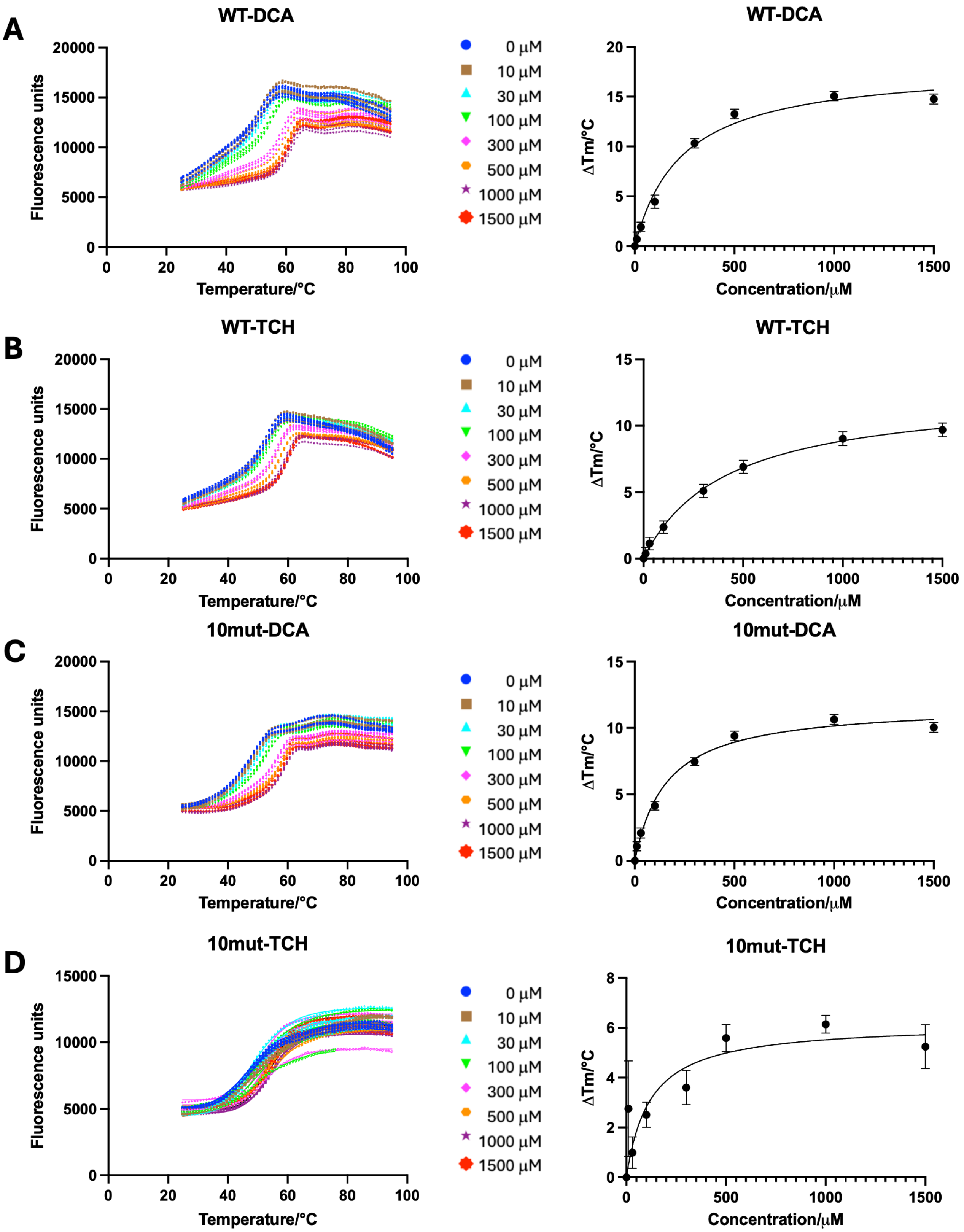


**Supplementary Figure 2: Stabilisation of $ASBT_{LB}$ by deoxycholate and taurocholate. A)** Wild type $ASBT_{LB}$ with DCA. The left panel shows the raw melting curves with increasing amounts of DCA with 4 technical repeats for each concentration. On the right panel, the ΔTM for each concentration is plotted as a function of concentration. Though stabilisation may be caused by multiple factors other than specific binding, the best fit through the points gives an apparent $K_d$ of 236 ± 40µM. **B)** Wild type $ASBT_{LB}$ with TCH ($K_d$ 416 ± 30µM). **C)** $ASBT_{LBmut}$ with DCA ($K_d$ 159 ± 25µM). **D)** $ASBT_{LBmut}$ with TCH ($K_d$ 119 ± 81µM).

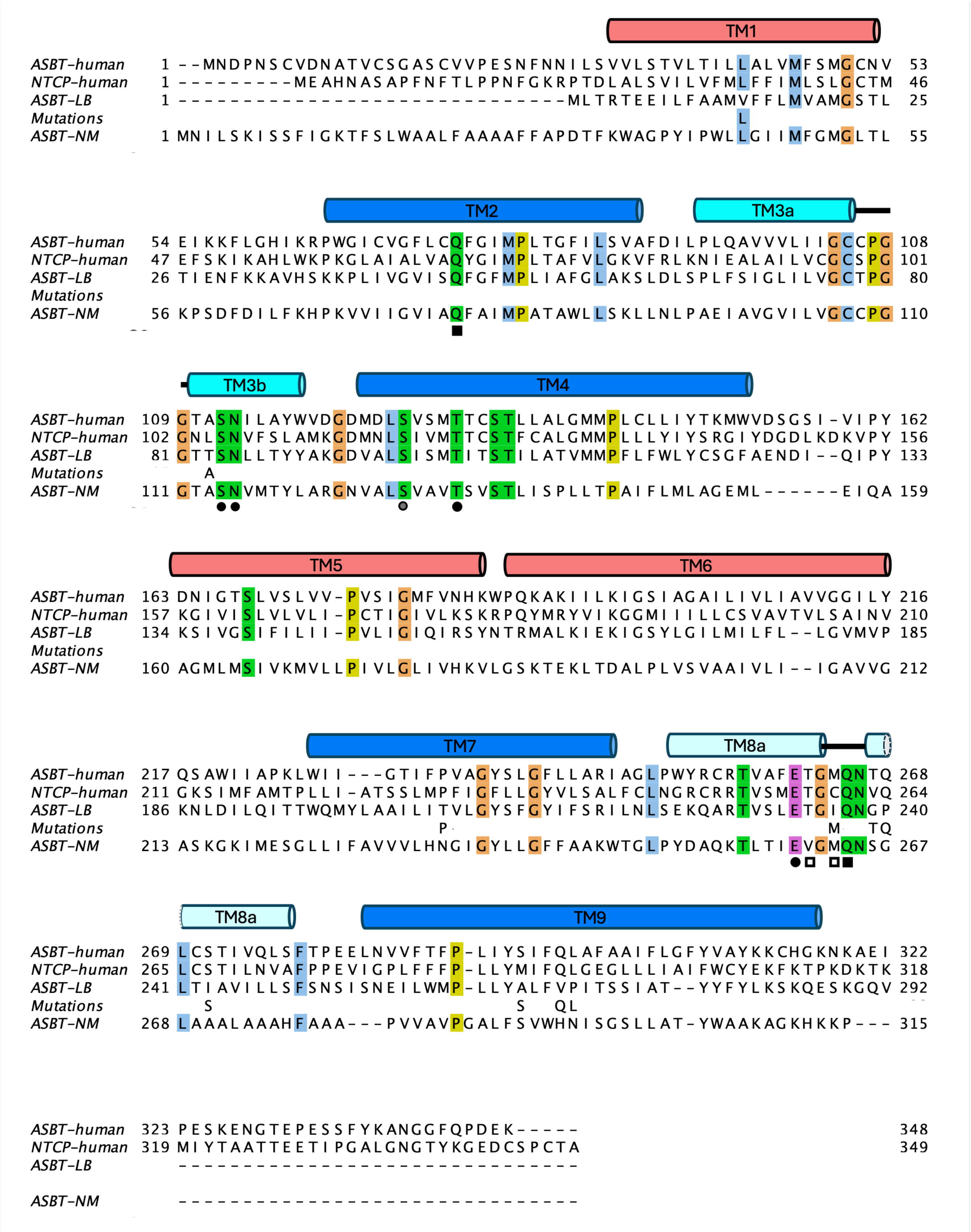


**Supplementary Figure 3: Sequence alignment of $ASBT_{LB}$ highlighting the mutations introduced to $ASBT_{LB}$.** The secondary structure and sodium-binding residues are shown as in Supplementary Figure 1. hNTCP (Q14973) and $ASBT_{NM}$(Q9K0A9) are also included.

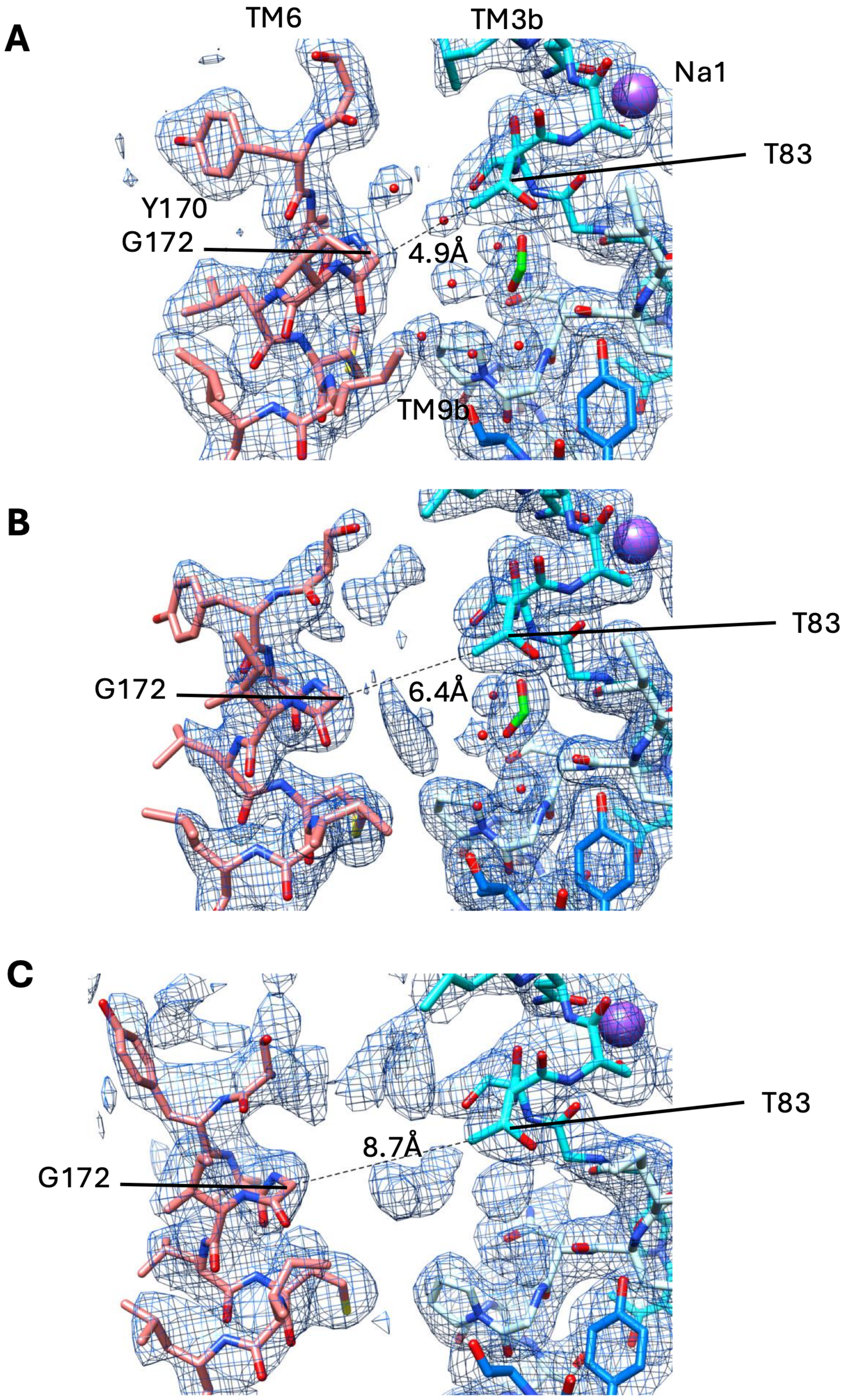


**Supplementary Figure 4: Electron density in the expected substrate binding site.** **A)** ASBT$_{LBWT\text{-}Xtal1A}$. Formate (green carbon atoms) from the crystallisation solution has been modelled. The dashed line indicates the distance between Thr 83 in the core domain and Gly 72 on TM6 of the panel domain. **B)** ASBT$_{LBWT\text{-}Xtal1B}$. Formate has also been modelled, but additional density, presumably lipids is also observed between the panel and core domains. **C)** ASBT$_{LBxtal2}$.

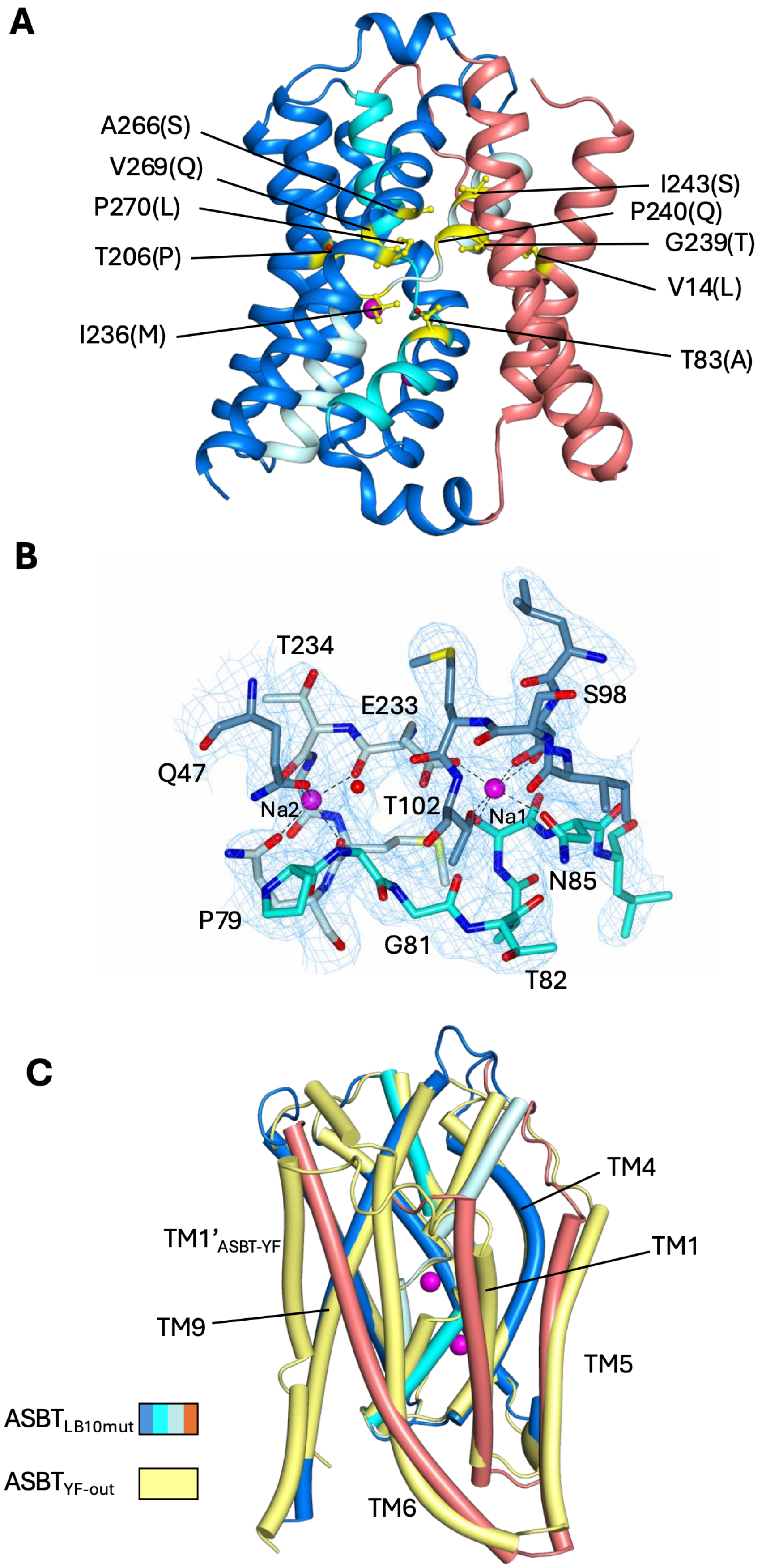


**Supplementary Figure 5: ASBT$_{LB10mut}$.** **A)** Location of mutations introduced ASBT$_{LB}$ to create ASBT$_{LBmut10}$. The mutations are indicated in yellow. **B)** Electron density associated with the Na binding sites for ASBT$_{LBmut10}$. The 2mFo-Fc density has been contoured at 1σ. **C)** Overlay of ASBT $_{LBmut10}$ with ASBT$_{YF}$ (4N7X; coloured yellow).

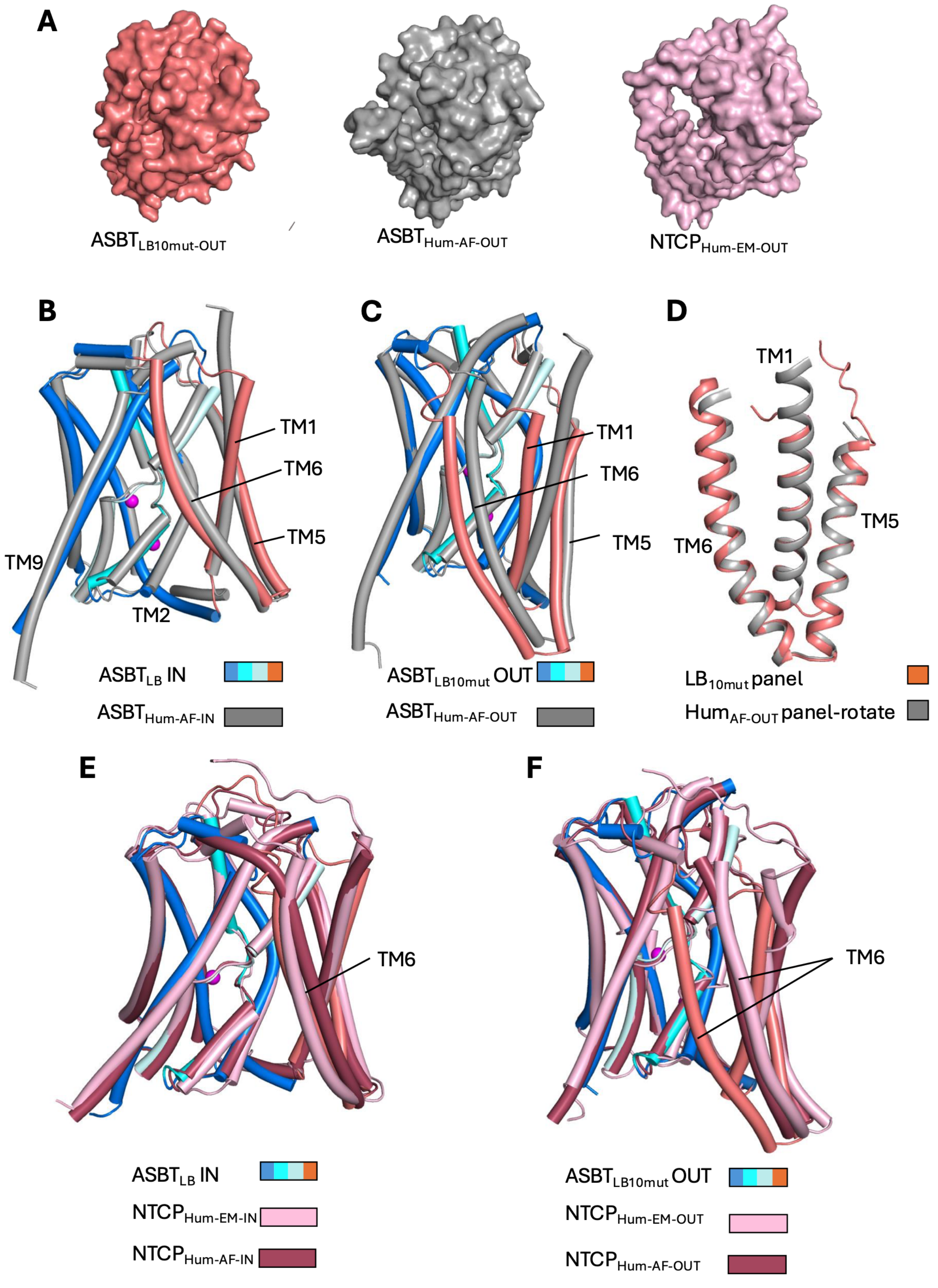


**Supplementary Figure 6: Comparison with structures calculated in AlphaFold2. A)** Surface representation of the outward-facing structures. **B)** Superposition of ASBT$_{LB\text{-}xtal1A}$ on a predicted inward-facing conformation of hASBT (grey). **C)** Superposition of ASBT$_{LBmut10}$ on a predicted outward-facing conformation of hASBT. **D)** Superposition of the respective panel domains from B. **E)** Superposition of ASBT$_{LB\text{-}xtal2}$ on inward-facing structure of hhNTCP (7PQG; coloured as in Figure 3) and an AlphaFold2 prediction (deep red). **F)** Superposition of ASBT$_{LBmut10}$ on an outward-facing structure of hNTCP (7PQQ) and an AlphaFold2 prediction.

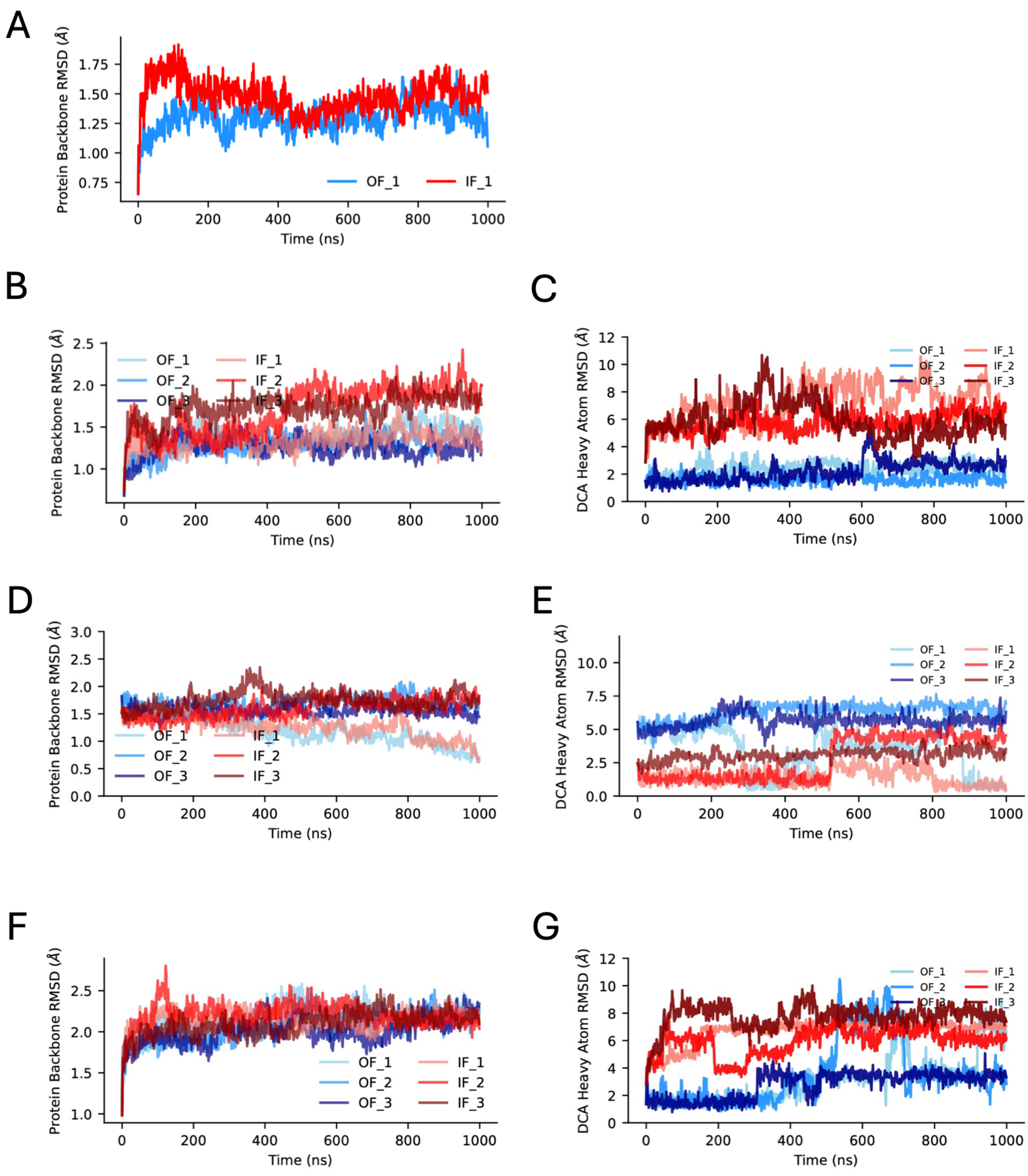


**Supplementary Figure 7: Conformational stability of MD simulations and DCA ligand binding as measured by RMSD after optimal structural superposition. A)** Protein backbone RMSD of apo WT $ASBT_{LB}$, after superposition on crystal structures (humanized $ASBT_{LB}$ for OF, WT $ASBT_{LB}$ for IF). **B)** humanized $ASBT_{LB}$ protein backbone RMSD (after superposition on the crystal structures) for three repeats with 2 $Na^+$, 1 DCA bound in OF (blue) and IF (red) conformation. **C)** Heavy atom RMSD of DCA bound to humanized $ASBT_{LB}$, after superposition of the protein on the DCA-bound crystal structure (for OF) or the protein with the initial Autodock/Vina docking pose (for IF). **D)** Protein backbone RMSD for WT with 2 $Na^+$/1 DCA. **E)** DCA heavy atom RMSD while bound to WT. **F)** Protein backbone RMSF for model of hASBT with 2 $Na^+$/1DCA. **G)** DCA heavy atom RMSD while bound to model of hASBT.

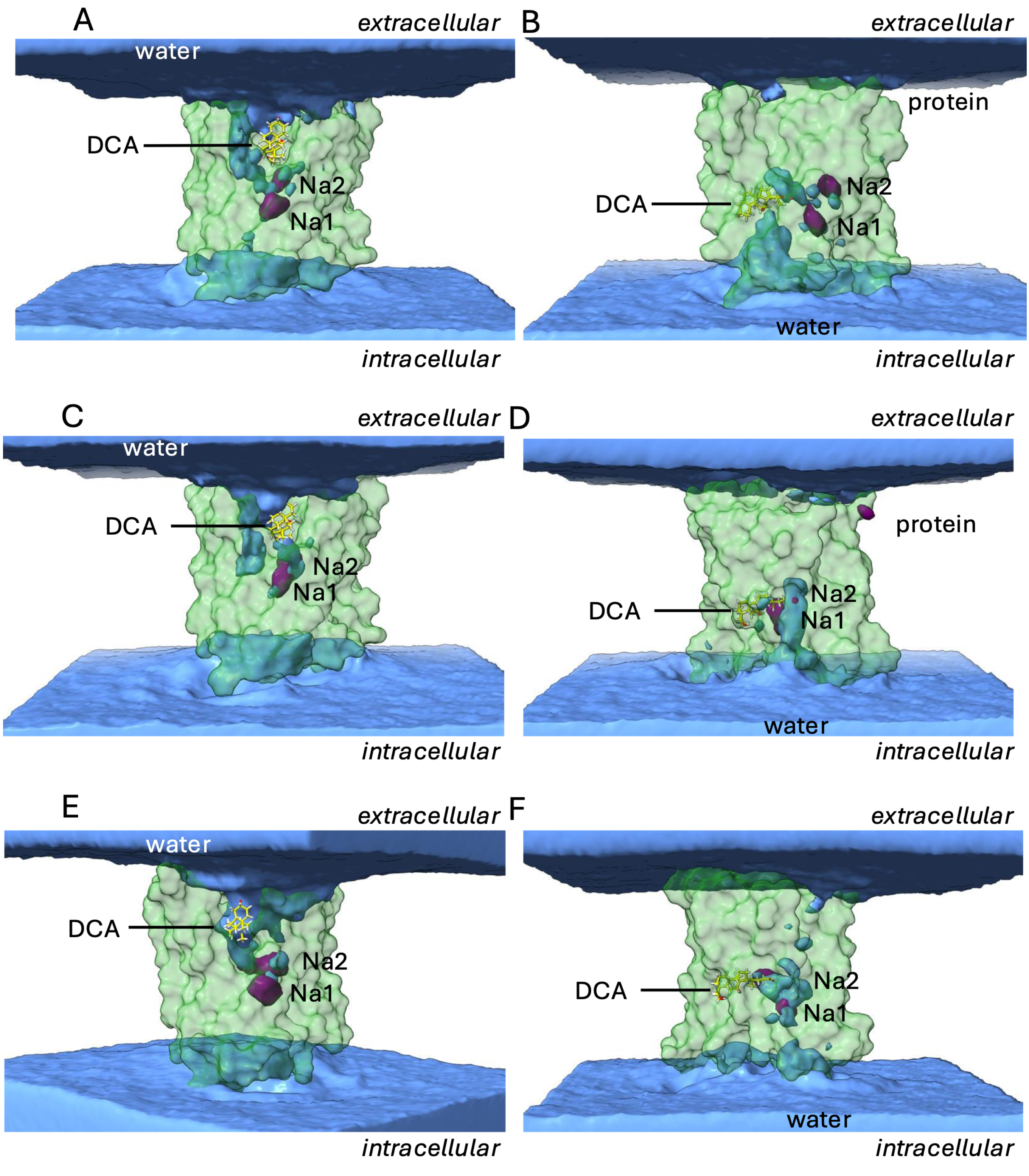


**Supplementary Figure 8: Accessibility of substrate and ion binding sites from MD simulations with 2 $Na^+$ and 1 DCA bound.** The density of water is shown in blue at a contour of 60% of the bulk density. The density of $Na^+$ ions (purple) is contoured at 4.2 M and clearly shows high density in the Na1 and Na2 sites. DCA is shown with a representative pose in stick representation. The protein surface is shown in transparent green. All densities were averaged across all three simulation repeats. **A)** Humanized $ASBT_{LB10mut}$ outward facing **B)** Humanized $ASBT_{LB10mut}$ inward facing. **C)** WT OF. **D)** WT IF. **E)** hASBT OF. **F)** hASBT IF.

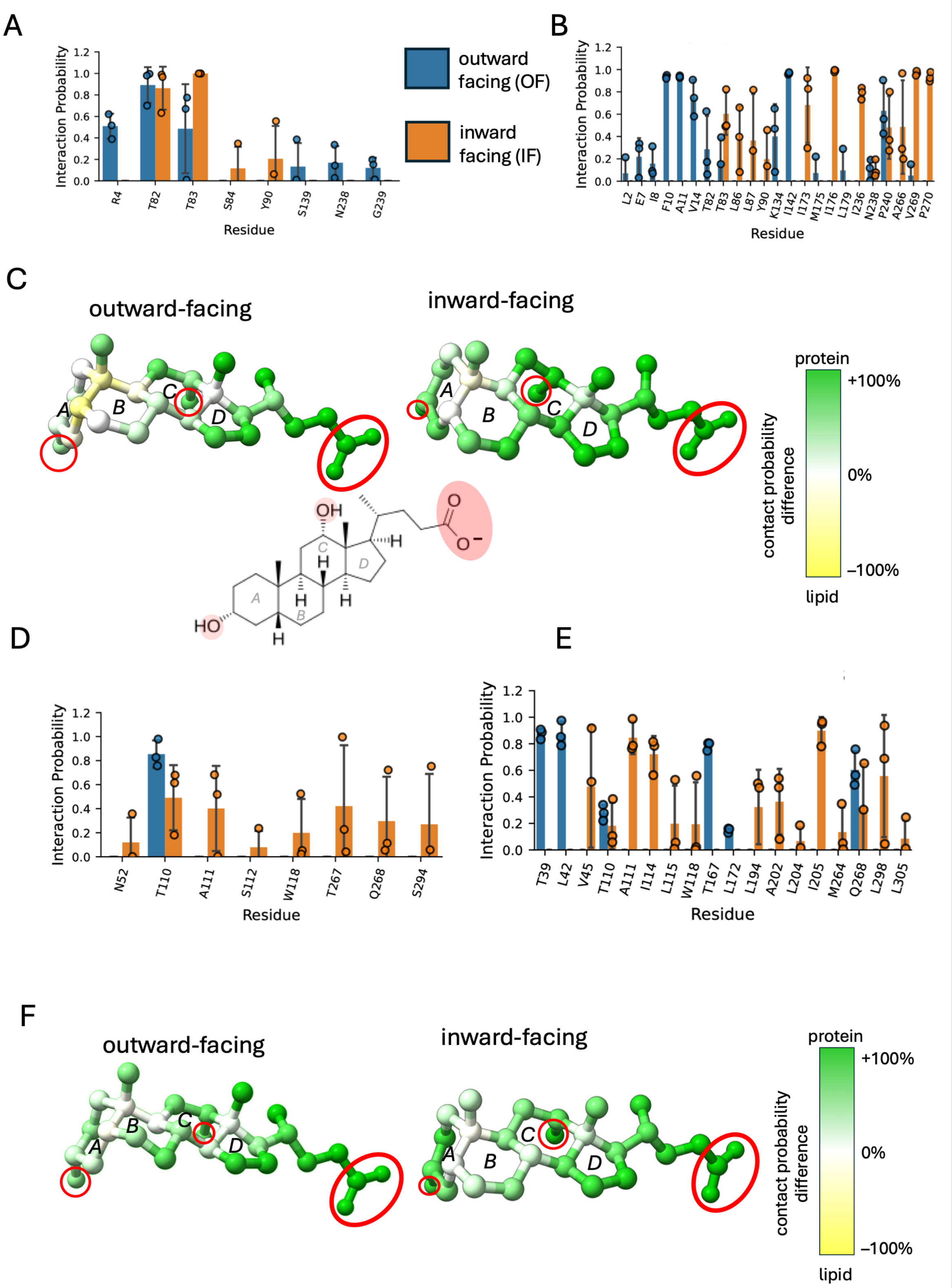


**Supplementary Figure 9**: **DCA binding in WT ASBT$_{LB}$ and hASBT. A)** Hydrogen bonding acceptor interaction probability between DCA and specific residues in WT ASBT$_{LB}$. **B)** Hydrophobic contact probability between DCA and specific residues in WT ASBT$_{LB}$. **C)** Difference of contact probability between DCA-protein and DCA-lipids, projected on the DCA structure, when bound to OF and IF WT ASBT$_{LB}$. Polar groups are marked red in DCA. **D)** Hydrogen bonding acceptor interaction probability between DCA and specific residues in hASBT. **E)** Hydrophobic contact probability between DCA and specific residues in hASBT. **F)** Difference of contact probability between DCA-protein and DCA-lipids, projected on the DCA structure, when bound to OF and IF WT hASBT.

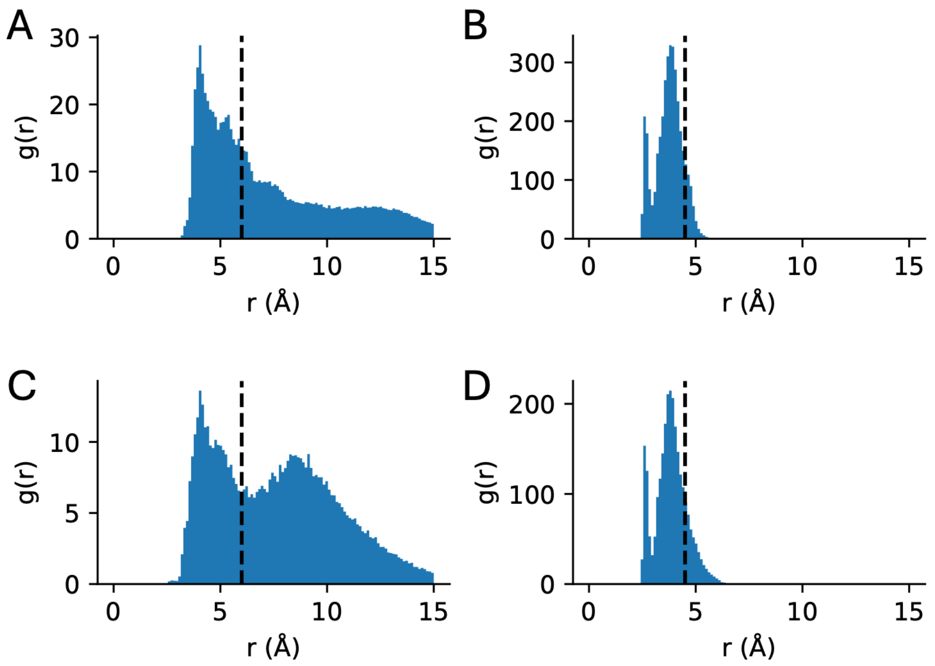


**Supplementary Figure 10:** *g(r)* radial distance distributions and chosen distance cutoffs (black line) for **A**. OF DCA-lipid, **B**. OF DCA-protein, **C**. IF DCA-lipid, **D**. IF DCA-protein interactions. All systems contained WT $ASBT_{LB}$. Cutoff distances were chosen manually to roughly approximate the boundary of the first interaction shell (DCA-protein 4.5 Å and DCA-lipids 6 Å).

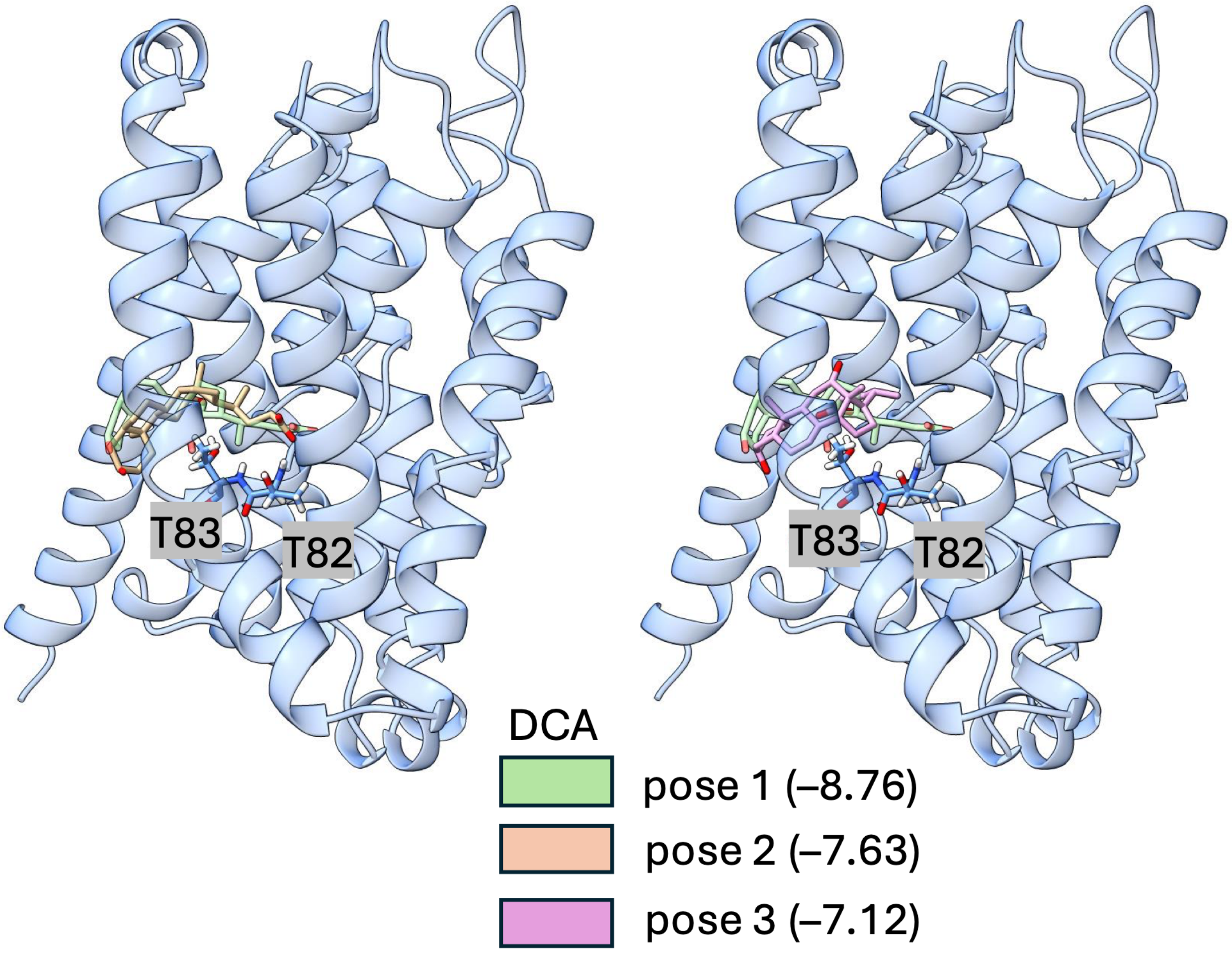


**Supplementary Figure 11:** Docking poses of DCA to the WT IF structure. The IF structure of $ASBT_{LB}$ is shown in light blue. The best scoring pose 1 (AutodockVina score -8.76) is shown in light green and compared to the second best pose 2 (score -7.63) in light brown and the third best pose 3 (score -7.12) in light purple. Residues Thr82 and Thr83 are highlighted.

**Supplementary Movie 1**

**Morph showing the conformational differences between the five structures of $ASBT_{LB}$.** The superpositions were made in ChimeraX and show the transition from $ASBT_{LB10mut}$ to $ASBT_{LB10mut\text{-}DCA}$ to $ASBT_{LBxtal2}$ to $ASBT_{LBxtal1b}$, to $ASBT_{LBxtal1a}$. The protein is coloured as in Figure 1.
